# Experimental exposure to winter thaws reveals tipping point in yellow birch bud mortality and phenology in the northern temperate forest of Québec, Canada

**DOI:** 10.1101/2023.10.20.563331

**Authors:** Benjamin Marquis, Geneviève Lajoie

## Abstract

Climate change is expected to increase the frequency and intensity of winter thaws, which could have two contrasting effects on leaf phenology. Phenology could either be advanced through the acceleration of forcing accumulation or chilling completion, or be postponed through a reduction in chilling associated with warming air temperature. We tested the influence of winter thaws on budburst phenology by exposing 300 tree cuttings of sugar maple and yellow birch trees to five different frequencies and durations of winter thaws in the lab. In spring, half of the cuttings were exposed to air temperature in two cities representing an air temperature gradient of + 2.0 °C to mimic the ongoing climate warming and bud phenology was monitored three times a week. Irrespective of thaw treatment, yellow birch phenology occurred earlier in the warmer city, showing the importance of spring temperature in triggering budburst. The treatment with the highest frequency and duration of thawing increased bud mortality and delayed the onset of spring phenology whereas low frequency treatments did not, thereby identifying a tipping point in the impact of winter thaws on bud phenology. Past this point, winter thaws could slow the acceleration of bud phenology induced by warmer spring temperature and limit carbon uptake by delaying the closure of the canopy. Climate change simulations projected by the CMIP6 Canadian downscaled climate scenario show that winter thaws will increase in frequency. Hence the expected advance in the spring phenology associated with warmer spring is not necessarily as straightforward as previously thought.

**Graphical abstract:** 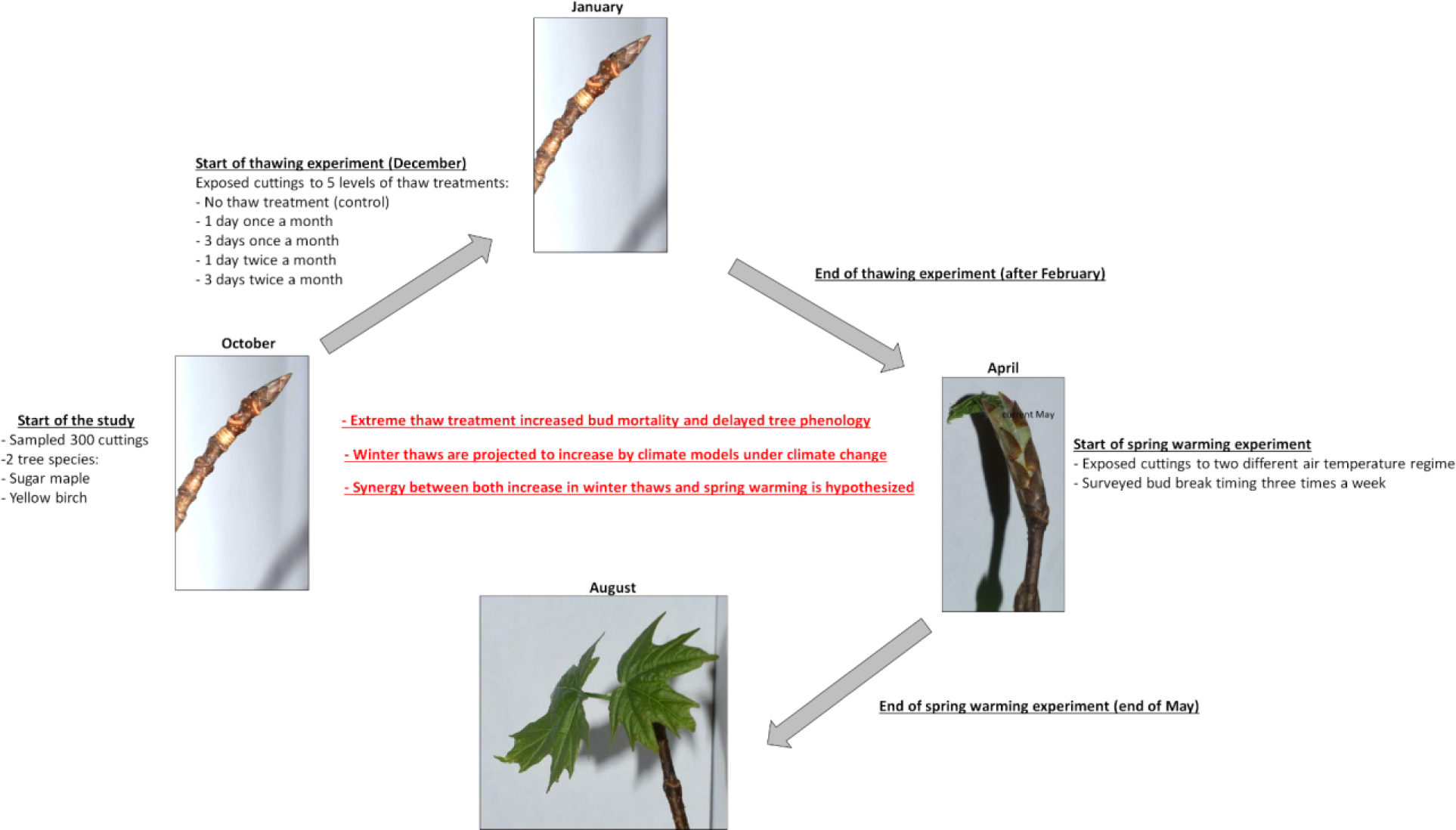

## 1.0 Introduction

The spring phenology of temperate and boreal tree species is advancing under climate change. While this shift in phenology could favor tree growth by extending their growing season, it could also be detrimental if it increases tree exposure to extreme climatic events such as droughts or spring frosts or if it increases susceptibility to leaf predation by insects [1–9]. Understanding the response of tree phenology to climate change is therefore key to predict how global processes such as carbon sequestration by forest ecosystems could be affected by climate change through their impacts on tree biomass and mortality [7, 10–14]. Indeed, shifts in the synchrony between air temperature and tree phenology can change the net carbon uptake by trees of the northeastern temperate forest by 4 g C m-2 per shifting day [15].

Bud phenology of temperate and boreal tree species is known to be driven by the dual impact of air temperature (chilling, and forcing) and their interactions [16,17]. Based on recent literature, chilling needs consist in air temperature in the range of −3 to 10 °C occurring from late summer to late winter that interacts with photoperiod to trigger the physiological processes of cold acclimation and dormancy release, whereas forcing needs? consists in heat that triggers the bursting of buds, calculated as growing degree-days based on the mean daily air temperature above zero starting on January [18–26]. According to the alternating model, increasing the number of chilling requirements decreases the forcing required for the bursting of buds [18,27]. In addition, abnormally high air temperatures late in the winter or early in spring can trigger the bursting of buds which can lead to false spring events [28].

One aspect of air temperature that has been overlooked in bud phenology models but could potentially impact tree phenology is winter thaws. Winter thaws are typically considered as being events during which air temperature goes above and below the freezing point during a day, also referred to as day with zero-crossing [29,30]. The increase in air temperature above zero for a few hours during the winter can be sufficient to reduce the cold hardiness of trees [31,32]. However, given air temperature remains above zero degrees only for a few hours, this forcing might not be captured by the growing degree-days calculated using the daily average air temperature. Therefore, how these thaw events impact tree phenology through changes in chilling and forcing accumulation needs to be elucidated. Two contrasting effects of winter thaws on bud phenology have been hypothesized. First, an increase in the frequency and duration of winter thaws could advance the spring phenology by accelerating forcing accumulation [33,34]. Second, a reduction in chilling accumulation linked with higher winter air temperature could delay the spring phenology. With the ongoing warming, winter thaws are expected to increase in frequency, thus, play a greater role in triggering bud break timing. However, climate change is not equal across biomes, therefore, the rate of chilling and forcing accumulation as well as the frequency (number of events), duration (number of days thaw event last) and intensity (air temperature reached during the thaw event) of winter thaws should also be site-specific, which prevents extrapolating results obtained on few species and locations to other species and sites and stresses the need to study many species and sites.

In eastern Canada, freeze-thaw cycles are most frequent between parallels 45° N and 47 °N (https://climatedata.ca/explore/variable/?coords=47.01797041194111,-68.63184644428942,6&delta=&dataset=cmip6&geo-select=&var=dlyfrzthw_tx0_tn-1&var-group=other&mora=ann&rcp=ssp126&decade=1990s&sector=), which encompasses most of the northern temperate forests composed of ecologically and economically important tree species such as sugar maple and yellow birch trees [35–37]. Therefore, understanding the response of tree phenology to the change in freeze-thaw events that is projected under climate change is essential to predict change in the functions of the temperate deciduous forest ecosystem.

In this study we evaluate how winter thaws are projected to change during the period 1950 to 2100 at our study site using twelve climate models from the Canadian downscaled climate scenarios (CMIP6). Then, we determined the effect of increasing frequency and duration of winter thaws on the bud phenology of sugar maple and yellow birch trees by experimentally exposing dormant tree cuttings to different levels of thaws during the winter, then exposing them to contrasting air temperature in spring representing the projected increase in spring temperature under the current pace of climate change. We first hypothesized that global warming will increase the frequency and duration of winter thaws. Concerning spring phenology, we tested two hypotheses, first, that increasing the forcing in spring would advance the onset of the bursting of buds irrespective of the thaw treatment. We thus predicted that cuttings at the warmer site would open their buds earlier compared to the colder site. Second, we tested the hypothesis that increasing the number and duration of winter thaws would advance the onset of the bursting of buds by either contributing to the chilling needs or the forcing in spring. Therefore, we predicted that the timing of bud burst would be correlated with the duration and frequency of the thaw treatments, with cuttings exposed to more thawing bursting first.

## 2.0 Materials and methods

### 2.1) Sampling site

Sample collection took place in the Mont-Orford National Park (−72.19 °W; 45.32 °N) which is located in the southern temperate forest of Québec and lies in the Sugar maple and basswood bioclimatic domain [38] (Fig. 1). This park was created in 1938 and covers an area of 59.5 km^2^ around the Mont-Orford Mountain, which is part of the Appalachian Mountain Chain. Mature sugar maple stands cover 75 % of the area of the park however the vegetation shifts towards conifer stands (e.g. balsam fir; *Abies balsamea*) at high elevation and eastern hemlock (*Tsuga canadensis*) at wetter low-elevation sites. Based on climate normals, the mean annual temperature for the period 1981-2010 was 5.6 °C (climate ID: 7024440, weather station: Magog). The average air temperature is below freezing during the months of December, January, February, and March. January is the coldest month with the average air temperature reaching −10 °C and with extreme daily minimum reaching −37 °C. The frost-free period lasts about 140 days from May 15^th^ to October 3^rd^. During this period, 3000 growing degree-days - based on the 0 °C threshold - accumulate. The warmest month is July, with air temperature reaching, on average, 19.4 °C with extreme daily maximum reaching 34.4 °C. Total precipitation sum is 1142 mm of which 903 mm falls as rain and 240 cm falls as snow.

**Fig. 1.**
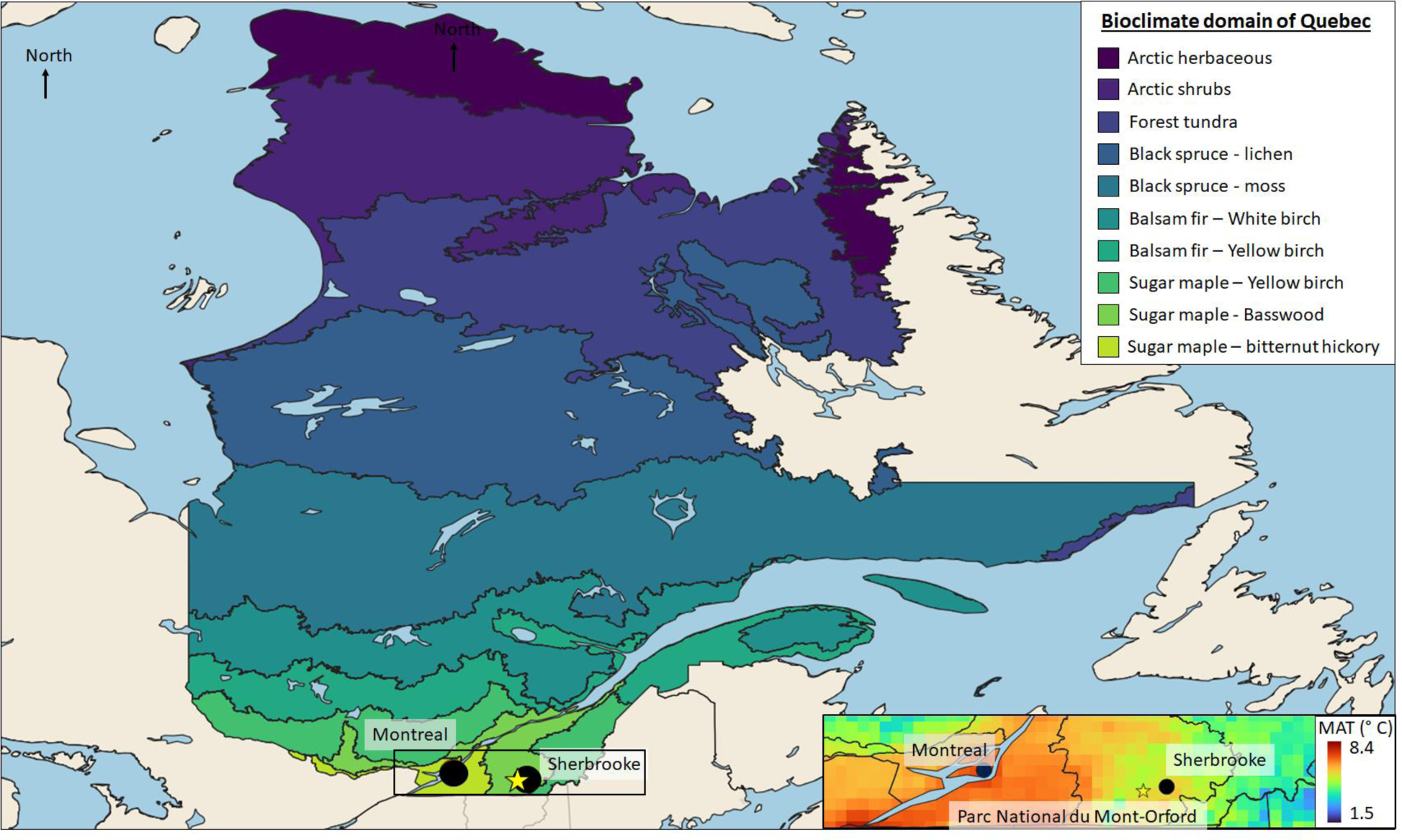
Map showing the bioclimate domains of Québec, the locations of the regrowth tests (black dots), the location of the national park where sample collection took place (the yellow star). The inset shows the average of the mean annual temperature around our study area for the period 1980-2020 simulated using the BioSim 11 software at a resolution of 0.05 degree and accounts for the effects of longitude, latitude, and elevation.

### 2.2) Projected winter thaw events under climate change

To calculate the change in frequency and duration of winter thaws under climate change we used the daily minimum and maximum air temperature that is projected by 12 chains of global and regional climate models that have a resolution of 10 km, cover the period 1950-2100, and run on 366 calendar days from the Canadian Downscaled Climate Scenarios (CMIP6) for two socioeconomic pathway (SSP2-4.5 and SSP5-8.5). Specifically, we analyzed the projections of the following climate models: ACCESS-CM2, ACCESS-ESM1-5, CNRM-CM6-1, CNRM-ESM2-1, EC-Earth3, EC-Earth3-Veg, IPSL-CM6A-LR, MIROC6, MIROC-ES2L, MPI-ESM1-2-HR, MPI-ESM1-2-LR, MRI-ESM2-0. The daily data were downscaled following the bias corrected constructed analogs [BCCA; 39] and the Quantile Delta Mapping [QDM; 40] and are freely available from the Pacific Climate Impacts Consortium (PAVICS) data portal (https://data.pacificclimate.org/portal/downscaled_cmip6/map/).

We analyzed the past and projected change in three complementary variables describing thawing frequency and duration: the frequency of daily winter thaws, the mean duration of long winter thaws and the number of thawing days. The frequency of daily winter thaw events was defined as the number of days when the maximum daily air temperature reaches above 0°C and the minimum air temperature reaches below 0 °C, hence, a day with zero crossing. The duration of long winter thaws were defined as the number of consecutive days during which the minimum air temperature stayed above 0 °C. The number of thawing days was calculated as the total number of days during which the max temperature was above 0°C, hence considers both daily thaws and each day of long winter thaws. For analyses, the number of daily thaws and the total number of thaw days were summed per winter months (January, February, March, and December) for each year whereas the duration, in days, of each long term thaw was averaged per month for each year when it was projected to occur. The temporal trends in each three complementary variables describing the thaw events were analyzed per winter months using linear regressions implemented in the R software for statistical computing (version 4.3.1). The average temporal trend of all climate models is presented in the result section 3.1 and the temporal trends for each climate model as presented in Figs S1-S12.

### 2.3) Field data and plant material collection

On October 29th, 2022, we identified fifteen dominant or codominant healthy sugar maple, *Acer saccharum,* trees (average DBH was 14 cm ranging from 7 cm to 32 cm) and fifteen yellow birch, *Betula alleghaniensis,* trees with their crown in the canopy (average DBH was 13.9 cm ranging from 6 cm to 29 cm) in a low elevation (285 - 290 m. above sea level) stand dominated by sugar maple and yellow birch trees and with beech (*Fagus grandifolia*) as the dominant understory tree. On each identified tree, we cut branches from the lower part of the crown (within the first 4 meters above the ground) to obtain ten cuttings of 20 cm with a dormant terminal bud. During field work, cuttings were kept in sealed plastic bags. Then, on seven randomly selected trees, we attached a thermometer (iButtons https://www.ibuttonlink.com/collections/thermochron/products/ds1921g) at breast height to record air temperature every 4 hours until spring 2023 (June 4). This sampling rate captured the daily variation in air temperature which allowed us to detect winter thaws at the field site and compare it to our thaw treatments and the thaws projected by the climate models. The *in situ* air temperature data recorded by each thermometer at the field site were then averaged to obtain a single daily record of air temperature. Back from the field, cuttings were kept in their plastic bags and placed in the dark at 4 °C in a refrigerator until December 1^st^ the date when they were transferred to a freezer set at −10 °C. On March 1^st^, cuttings were transferred back to the refrigerator (4 °C). These transitions from cold (4 °C) to freezing air temperature (−10 °C) and back to cold (4 °C) air temperature allowed cuttings to acclimate and de-acclimate to freezing temperatures as well as to fulfill chilling requirements, since air temperature around 4 °C contribute the most to chilling accumulation [19, 27, 34]. Thereby, we limited the potential effect of the lack of chilling on the subsequent timing of budbreak in spring [41]. By collecting ten cuttings from the same tree and submitting them to different winter thaw treatments we could also control for potential genotype effects.

### 2.4) Winter thaw treatments

To determine the response of trees to winter thaws we submitted cuttings to different frequencies and durations of winter thaws. Thaws were mimicked by pulling the cuttings out of the freezer and exposing them to an air temperature of 4 °C, either for 24 (low duration) or 72 (high duration) hours. These thaw treatments were conducted either once a month (low frequency of winter thaws) or twice a month (high frequency of winter thaws) from early December to early March, including a no-thaw control. We therefore used a set of five different thawing treatments for each of two tree species to determine the response of bud phenology to winter thaws (Table S1). Since we wanted to test the effect of air temperature only, our thaw treatments were conducted in the dark. On March 16th, cuttings were all removed from the freezer and kept at 4 °C for three weeks until the regrowth tests. To compare if our thaw treatments did represent the projected frequency and duration of future thaw events, we compared our thaw treatments to the projected thaw events during the period 1991-2020 and the period 2071-2100. We also compared it to *in situ* thaw events registered at our field site (Mont-Orford National Park).

### 2.5) Regrowth tests

On April 8, the cuttings were put in 500-mL glass containers half filled with water and placed outside, evenly split, in each of two cities representing current air temperatures at the field site (Sherbrooke, 22 km east from the field site, see Fig. 1) and warmer air temperatures expected in the future from climate change (Montreal, 124 km to the west from the field site, see Fig. 1). Indeed, the mean annual temperature in Sherbrooke city is 5.6 °C, the total precipitation sum is 1145 mm, the average length of the frost-free season is about 120 days during which 3052 growing degree-days accumulates, it is thus very similar to the field site (Table 1 and section 2.1 *Sampling site*). At the warmer site, which is located in Montreal city the mean annual air temperature increases by 1.2 °C reaching 6.8 °C, the frost-free period is 25 days longer (lasting 165 days), starting 16 days earlier (on April 29) and ending 10 days later (on October 12) compared to the field site. During this frost-free period, 349 more growing degree-days accumulate, and the daily maximum air temperature can reach 37.6 °C compared to 34.4 °C at the field site (Table 1). It is therefore a suitable site to test for the potential effect of the warmer weather on timing of the bursting of buds. Also, by setting our experiment outside, we increased the realism of our study compared to maintaining cuttings in growth chambers with no daily cycle in air temperature or with a constant rate at which air temperature is changed. The dates at which phenological stages were reached in the field and in the experiment were similar (Fig. S13), suggesting that our experimental system is indeed representative of natural processes. We registered *in situ* air temperature by setting four thermometers (iButtons) at both sites set to record air temperature every 4 hours. The *in situ* air temperature data recorded by each thermometer were then averaged to obtain a single record of air temperature per city. Given there is, on average, a difference of only 1 minute in photoperiod between Sherbrooke and Montreal, we expect that air temperature will be the major variable affecting bud phenology compared to photoperiod. Water inside the glass containers was periodically changed to maintain cleanliness.

**Table 1.** Climate normal from 1981-2010 at our field and experimental sites.

| Sites | MAT<br>(°C) | Growing<br>degree-days | Frost-free period |  |  | Daily<br>minimum<br>temp.<br>(°C) | Daily<br>maximum<br>temp. (°C) | Total<br>precipitation<br>sum (mm) | Photoperiod<br>(hours) | Weather<br>station<br>ID |
| --- | --- | --- | --- | --- | --- | --- | --- | --- | --- | --- |
|  |  |  | <u>Duration</u> | <u>Start</u> | <u>End</u> |  |  |  |  |  |
| Field | 5.6 | 3000 | 140 | May<br>15<sup>th</sup> | October<br>3<sup>rd</sup> | -38 | 34 | 1142 | 12.2 | 7024440 |
| Sherbrooke | 5.6 | 3052 | 120 | May<br>24<sup>th</sup> | September<br>23<sup>rd</sup> | -42 | 35 | 1145 | 12.2 | 7020860 |
| Montreal | 6.8 | 3349 | 165 | April<br>29<sup>th</sup> | October<br>12<sup>th</sup> | -37 | 37 | 1000 | 12.2 | 7025250 |

### 2.6) Bud phenology observations

For the regrowth experiments, the phenological stage of terminal buds was assessed three times per week starting on April 10th, two days after the transfer of cuttings from the refrigerator (4 °C) to the outside, in both Sherbrooke and Montreal. Monitoring of individual branches stopped when their terminal leaves had fully expanded or if the bud had died. The last observation was made on May 26th. Changes in bud phenology from dormant buds to fully open leaves was determined following a categorical scale comprising 7 stages [42]: dormant bud, initial bud swelling, bud elongation, bud swelling/green tip, bud break, extended bud break, initial leaf emergence and leaves fully expanded. However, due to difficulty in discerning stages one and two, both stages were merged for further analyses. Bud phenology of the 30 identified trees in the field was also monitored once a week starting on April 29 until leaves were fully expanded.

### 2.7) Statistical analyses of experimental data

We analyzed bud break phenology based on the day of the year (DOY) at which buds transitioned from one category to the next. Statistical analyses focused on three stages: the onset of budbreak (stage 1), the day at which bud break happened (stage 4), and the day at which leaves were fully expanded (stage 7). We estimated transition dates that occurred in-between two observation dates (i.e. that were not observed for a given stage) by interpolating the data linearly. Given observations were conducted three times a week, this interpolation added a maximum of two days per date, maintaining a high level of precision in the estimation of the timing of phenological development.

Thawing and spring warming treatments were first defined as categorical variables, respectively using the different thawing regimes and the cities as levels. Their effects on bud break were also calculated through their impact on the growing degree days accumulated by the cuttings (1) in each treatment between December 1^st^ (first day in the freezer) and April 8th (first day outside) for the thawing treatments, and (2) in each city between April 8th (first day outside) and the end of phenology observations (May 26th) for the spring warming treatment. The number of growing degree days accumulated during the thawing treatments was calculated as the cumulative sum of days during which each cutting was exposed to a temperature of 4 °C, multiplied by four (effect of thawing). To this sum we added the cumulative sum of mean daily temperatures above zero experienced in the city in which the cutting was placed outside to account for the effect of spring warming.

We first used mixed linear models to evaluate the impact of the thawing treatment, the spring warming treatment and their interaction on stages 1, 4 and 7 transition dates, using the tree as a random variable. We then evaluated their impacts through growing degree days accumulation by adding GDD accumulation and its interaction with thawing treatment and with spring warming treatment (city) to the model. We assessed if adding these terms to the model significantly increased the fit of the model using AIC. For stages 4 and 7, treatment 11 was not included in the models because less than three cuttings reached stage 4 in Montreal and less than three cuttings reached stage 7 in either city. To still evaluate the impact of treatment 11 on stage 4 phenology, We tested the influence of treatment and spring warming treatment on the proportion of cuttings reaching phenological stage 1, 4 and 7 using general linear models with a binomial error distribution. For stage 1, proportions of cuttings reaching the stage were calculated out of the total number of cuttings for each treatment and city combination (n=15). For stages 4 and 7, the number of cuttings reaching that stage were calculated out of the pool of cuttings that had respectively reached stage 1 and stage 4. For all phenological analyses, the species effects could not be measured provided the failure of most sugar maple buds to reach bud burst stage (stage 4). These analyses therefore focused on yellow birch data only.

## 3.0 Results

### 3.1) Temporal change in projected thaw events under climate change

We show that by 2100, daily thaws when air temperature rises and drops below zero within a single day are projected to increase, on average, by six, five, three, and five days during the winter months of January, February, March and December respectively under the socioeconomic pathways SSP-2.4.6 and by eight, nine, two, and six days under the most extreme warming scenario (SSP-5.85) compared to 1950 (Fig. 2). Daily thaws during March only marginally increased because air temperature likely warmed enough to cause prolonged thaw periods, thus when all days with a thaw event were analyzed, thaw events during the month of March are also projected to increase at a similar rate as during the other winter months (Fig. 2). Overall, by 2100, winters are projected to have, on average, 41 more thaw days compared to during 1950.

**Fig. 2.**
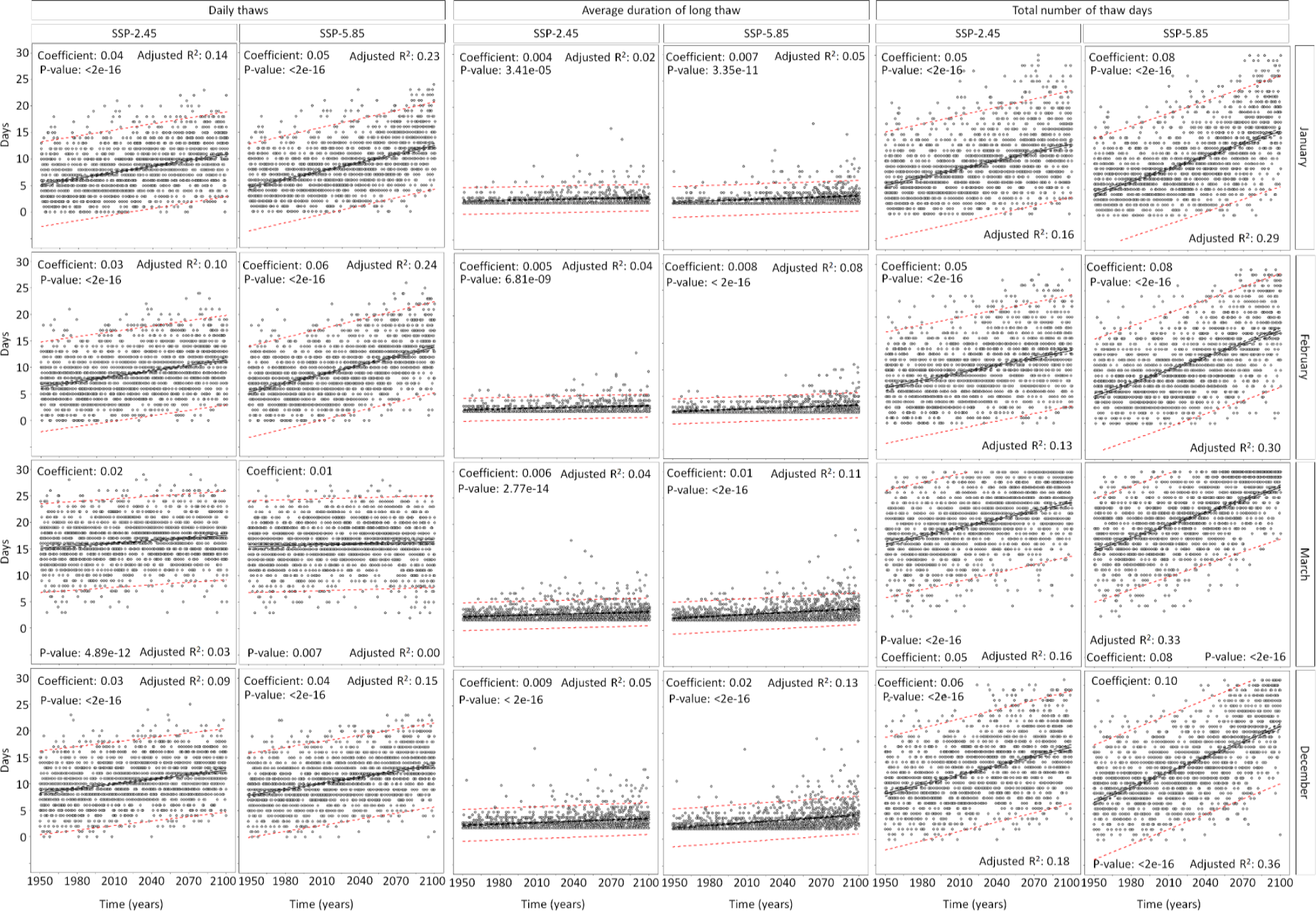
Temporal trends in the frequency of daily thaws, the average duration of a long thaw event (minimum air temperature > 0 °C) and finally the total number of days with a thaw events (daily + long term thaws) projected per winter months for each of the 12 climate models under two socioeconomic pathways. The plain black line represents the linear trend, the dashed black lines represent the 95 % confidence interval and the dashed red lines represent the prediction interval. Summary of the linear trends are reported in the figures.

We also report important inter-climate model variations in the rate at which thaw events are projected to increase for each winter month and under both socio economic pathways (Figs. S1-S12). The inter-climate model variability in the strength of the coefficient of the temporal trends of all thaw events (days when the maximum air temperature rises above zero) varied between 0.02 to 0.13 thaw/years in January, between 0.02 to 0.11 in February, between 0.03 to 0.10 in March and between 0.03 to 0.10 in December which resulted in an increase in 17, 14, 11, and 11 daily thaws between the climate model showing the strongest temporal trends and the climate model showing the smallest temporal trend. Under the socio-economic pathway 5.85 projecting extreme warming, the projected temporal trends in daily thaws varied from 0.03 to 0.14 in January, from 0.03 to 0.14 in February, from 0.05 to 0.11 in March and from 0.04 to 0.15 in December which resulted in an increase of 17, 17, 9, 17 daily thaws between climate models at both extremes of the spread. The adjusted R^2^ changed on average by + 13 % (ranging from + 4 to + 25 %), +18 % (ranging from + 4 to 26 %), + 16 % (ranging from + 3 to + 25), and + 19 % (ranging from + 3 to + 38 %) from SSP 2.45 to SSP 5.85 in January, February, March and December respectively. In general, under the socioeconomic pathway 2.46, the climate model that predicts the strongest temporal trends is ACCESS-CM2 except during the month March for which it is the climate model MIROC_ES2L whereas the climate model predicting the lowest temporal trends is ACCESS-ESM1-5. While the climate model ACCESS-ESM1-5 still projects the lowest temporal trends in thaw events under the socioeconomic pathway 5.85, the climate model predicting the strongest trends varied per month, being ACCESS-CM2 for the months of January and February, MIROC_ES2L for the month of March and EC_Earth3_Veg for the month of December.

### 3.2) Winter thaws comparison between climate models, in situ measurements and experimental treatments

By comparing the frequency of daily thaws and the duration, in days, of long term thaws, between the climate models, *in situ* thaw measurements and the experimental treatments, we realized that daily thaws are more frequent than we expected. Our most extreme treatment (three consecutive thaw days twice per month) exposing cuttings to six thaw days per month was similar to the number of daily thaws projected by climate models for the socioeconomic pathway 2.46 and 5.85 during the period 1991-2020 but was lower compared to the projected daily thaws during the period 2071-2100 under both socioeconomic pathways and was also lower compared to the *in situ* daily thaw events (Table 2). However, the average duration of a long term thaw event, when the minimum air temperature remains above zero for a few days, was rarely projected by the climate models to exceed three days, and that under both socioeconomic pathways and for both periods (1991-2020; 2071-2100). In addition, long term thaw events were not projected to occur every year but their probability of annual occurrence increased from the period 1991-2020 to the period 2071-2100 and also increased under the socioeconomic pathway projecting the most warming (SSP-5.85) compared to the socioeconomic pathway expecting milder warming (SSP-2.46). Moreover, long term thaw events were only registered once in December and once in January at the study site and lasted 2 and 3 days respectively (Fig. S14). Hence, our thaw treatments did represent extreme but plausible duration of thaws. Similarly, the spring warming treatments that we used in the experiment were representative of climate change scenarios expected for our field site. Specifically, using the city of Montreal as a space-for-time substitution for future air temperature at the field sampling site was appropriate provided air temperature was higher by an average of + 2 °C in Montreal (+ 15 °C) compared to Sherbrooke (+ 13 °C) over the period monitored (Fig. S14).

**Table 2.**
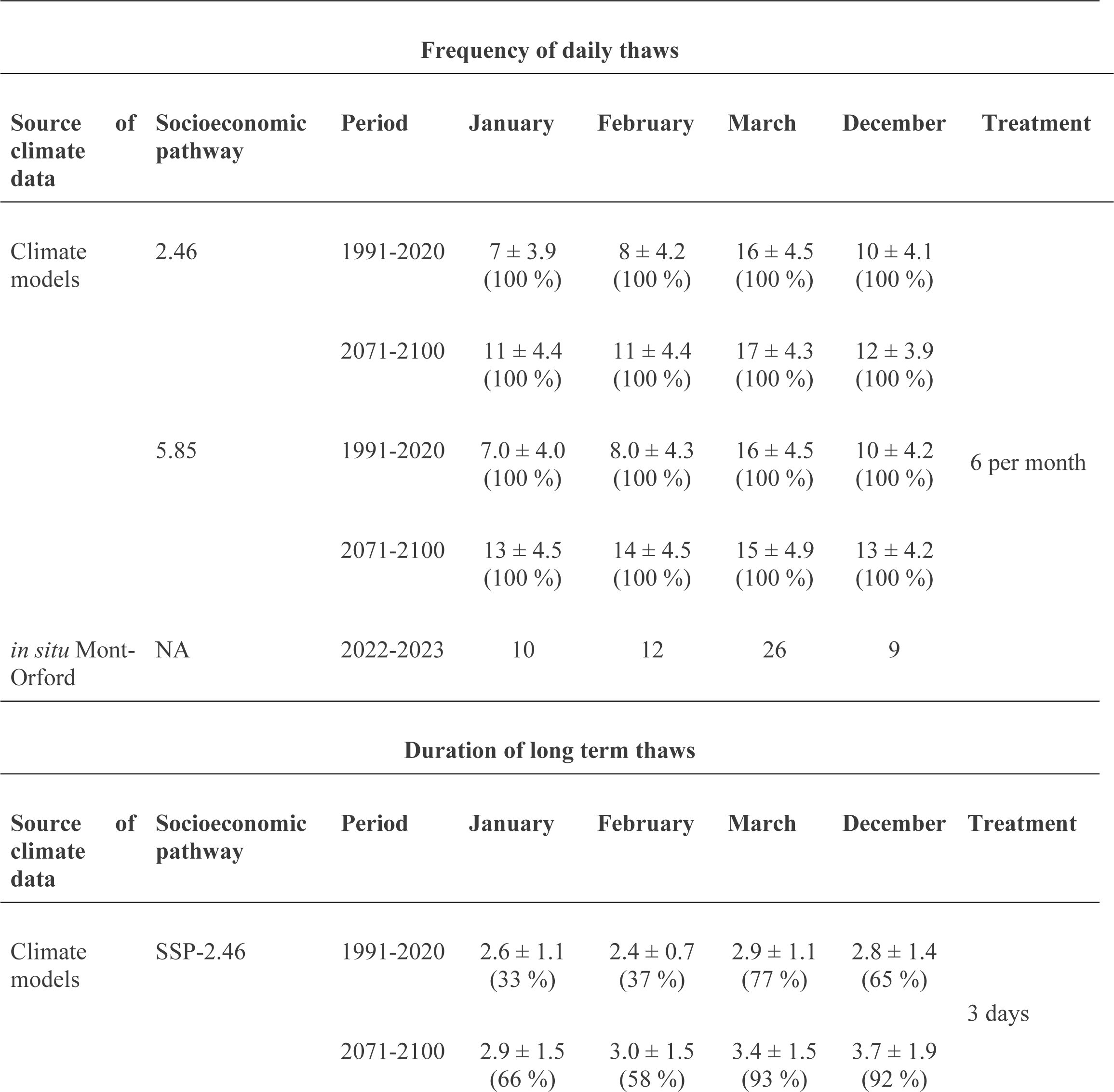
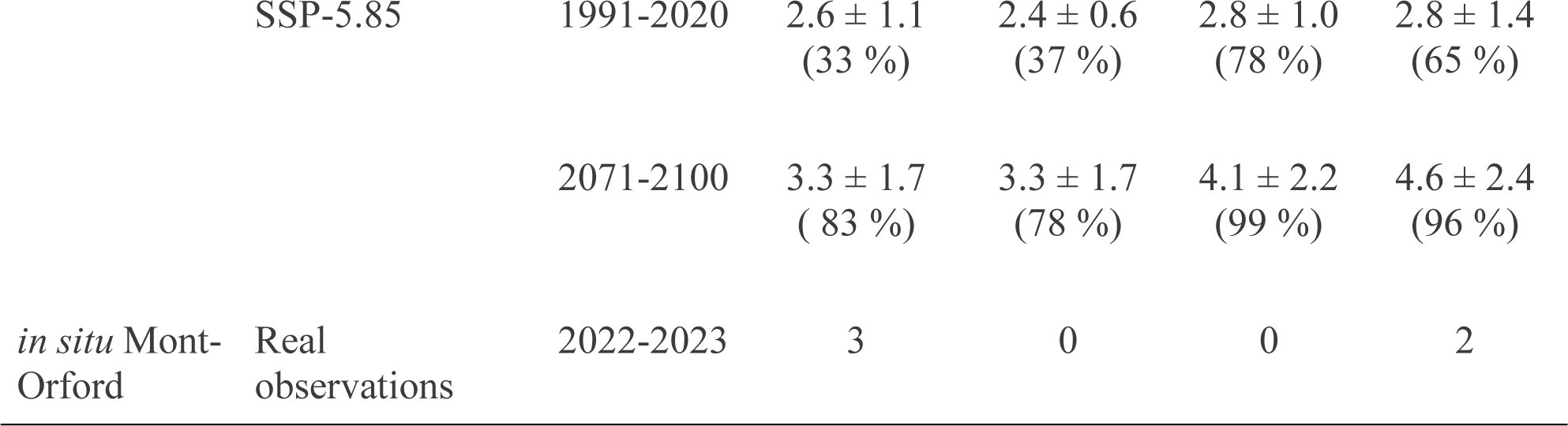
Comparison of the number of daily thaws, the duration, in days, of long-term thaws and their annual frequency (in %) between the climate models for current and future periods and two socioeconomic pathways, the thaw events registered *in situ* at the Mont-Orford National Park, and the thaw treatments. The annual frequency is reported in parenthesis and represents the number of times a daily thaw or a long thaw was projected by the 12 climate models.

### 3.3) Effect of winter thaws and spring warming on bud phenology

Winter thaw treatments and the amount of spring warming to which cuttings were exposed (i.e. the city effect) both impacted bud phenology. Winter thaw treatments were most important in explaining phenological development at early phenological stages (stage 1 and 4) (Fig. 3, Table 3). This effect was mostly driven by a delay in phenological development for cuttings that were in the most intense thawing treatment (treatment 11: three-day thaws, twice a month). This delay was observed at stage 1 for both cities and at stage 4 for the coldest city (Sherbrooke) (Fig. S15, S16). It could however not be measured in Montreal at stage 4, and for both cities at stage 7 provided too few cuttings from treatment 11 were able to pursue their phenological development across the season and ever reach those stages (Fig. 4). Indeed, the number of cuttings reaching phenological stages 4 and 7 were the lowest for the treatment with the longest duration of thaws (treatment 11: Table 4). Most thawing treatments however had a lower proportion of cuttings reaching stage 4 than the no-thaw control (Table 4).

**Fig. 3.**
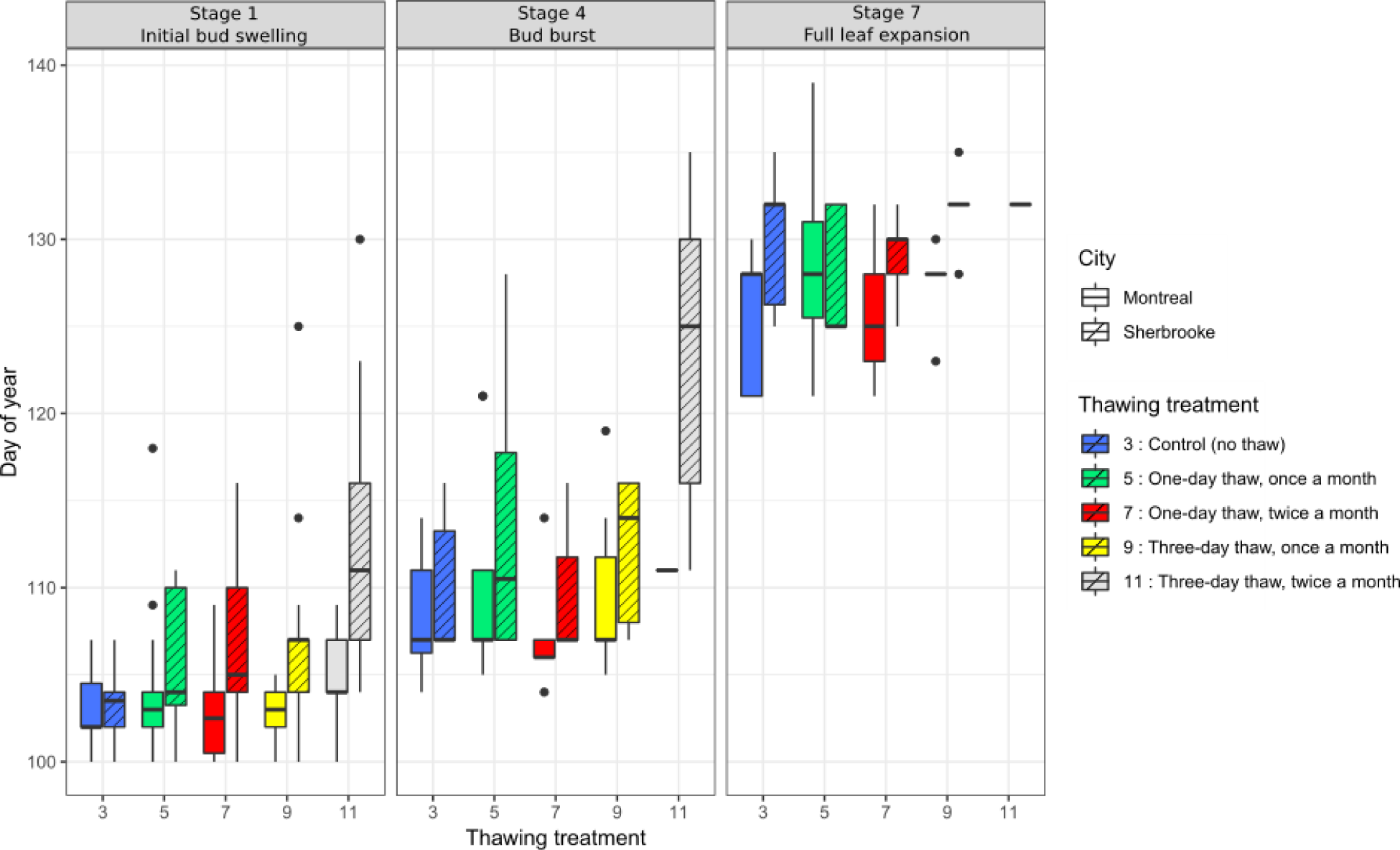
Day of year at which branches reach phenological stages 1, 4 and 7 by thawing treatment and city.

**Table 3.** Effect of thawing treatment, spring warming treatment (city) and growing degree days (GDD) on dates at which stage 1 (onset of budburst), stage 4 (bud break) and stage 7 (leaf fully expanded) phenological stages are reached by yellow birch cuttings. Comparisons between models not accounting for GDD (Model 0) and accounting for it (Model 1) are provided. Significance levels for each variable are indicated by * P < 0.05; ** P< 0.01; *** P < 0.001.

| **Stage 1: Initial bud swelling** |  |  |  |
| --- | --- | --- | --- |
| **i) Model 0** |  |  |  |
| **Factor** | **df** | **$\chi^2$ statistic** | **Pr(>Chisq)** |
| Treatment | 4 | 30.06 | <0.001 *** |
| City | 1 | 31.22 | <0.001 *** |
| Treatment:City | 4 | 18.35 | 0.001 ** |
| **ii) Model 1** |  |  |  |
| **Factor** | **df** | **$\chi^2$ statistic** | **Pr(>Chisq)** |
| Treatment | 4 | 729.06 | <0.001 *** |
| City | 1 | 63.55 | <0.001 *** |
| GDD | 1 | 3572.28 | <0.001 *** |
| Treatment:City | 4 | 14.02 | 0.007 ** |
| Treatment:GDD | 4 | 33.47 | <0.001 *** |
| City:GDD | 1 | 43.87 | <0.001 *** |
| **iii) Model comparison** |  |  |  |
| **Model** | **AIC** | **Chisq** | **Pr(>Chisq)** |
| Model 0 | 815.29 |  |  |
| Model 1 | 341.280 |  |  |
| Diff. | -474.01 | 486.01 | <0.001 |
| **Stage 4: Bud burst** |  |  |  |
| **i) Model 0** |  |  |  |
| **Factor** | **df** | **$\chi^2$ statistic** | **Pr(>Chisq)** |
| Treatment | 3 | 9.36 | 0.02 * |
| City | 1 | 4.84 | 0.03 * |
| Treatment:City | 3 | 0.64 | 0.88 |
| **ii) Model 1** |  |  |  |
| **Factor** | **df** | **$\chi^2$ statistic** | **Pr(>Chisq)** |
| Treatment | 3 | 1080.51 | <0.001 *** |
| City | 1 | 368.86 | <0.001 *** |
| GDD | 1 | 5080.31 | <0.001 *** |
| Treatment:City | 3 | 12.05 | 0.007 ** |
| Treatment:GDD | 3 | 10.03 | 0.02 * |
| City:GDD | 1 | 93.54 | <0.001 *** |
| **iii) Model comparison** |  |  |  |
| **Model** | **AIC** | **Chisq** | **Pr(>Chisq)** |
| Model 0 | 513.28 |  |  |
| Model 1 | 156.150 |  |  |
| Diff. | -357.13 | 367.13 | <0.001 |
| **Stage 7: Fully expanded leaves** |  |  |  |
| **i) Model 0** |  |  |  |

| Factor | df | $\chi^2$ statistic | Pr(>Chisq) |
| --- | --- | --- | --- |
| Treatment | 3 | 4.12 | 0.25 |
| City | 1 | 10.70 | 0.001 ** |
| Treatment:City | 3 | 4.58 | 0.21 |

**ii) Model 1**
| Factor | df | $\chi^2$ statistic | Pr(>Chisq) |
| --- | --- | --- | --- |
| Treatment | 3 | 663.87 | <0.001 *** |
| City | 1 | 1429.37 | <0.001 *** |
| GDD | 1 | 5757.56 | <0.001 *** |
| Treatment:City | 3 | 2.07 | 0.56 |
| Treatment:GDD | 3 | 7.07 | 0.07 |
| City:GDD | 1 | 3.68 | 0.05 |

**iii) Model comparison**
| Model | AIC | Chisq | Pr(>Chisq) |
| --- | --- | --- | --- |
| Model 0 | 339.93 |  |  |
| Model 1 | 60.87 |  |  |
| Diff. | -279.06 | 289.05 | <0.001 *** |

**Fig.4.**
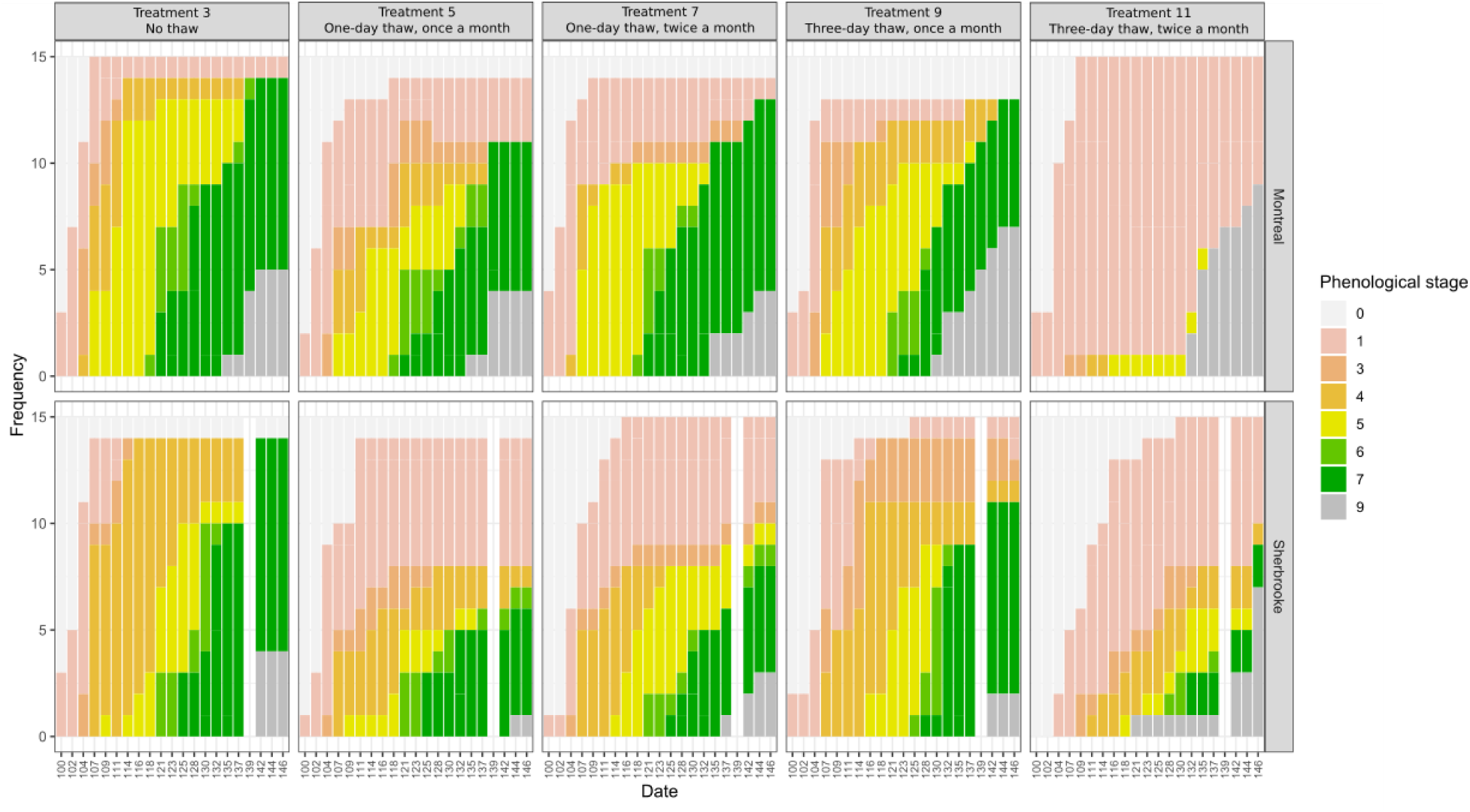
Proportion of cuttings at each phenological stage from day 100 to day 146 per treatment and city. Stage 9 corresponds to dead cuttings.

**Table 4.** Effect of thawing treatment and spring warming on the proportion of cuttings reaching phenological stages 1 (onset of budburst), 4 (bud break) and 7 (leaf fully expanded) across the duration of the experiment. General linear models with a binomial error distribution were run separately for each phenological stage. For stages 4 and 7, the number of cuttings reaching that stage were calculated out of the pool of cuttings that had respectively reached stage 1 and stage 4. Significance levels for each variable are indicated by * P < 0.05; ** P< 0.01; *** P < 0.001.

| **Stage 1: Initial bud swelling** |  |  |
| --- | --- | --- |
| **Variable** | **Coefficient (Std. Error)** |  |
| Intercept | 2.954 (1.038) | ** |
| Treatment 5 | -1.794 (1.138) |  |
| Treatment 7 | 0.000 (1.444) |  |
| Treatment 9 | -1.188 (1.193) |  |
| Treatment 11 | 0.000 (1.444) |  |
| City Sherbrooke | 1.097 (0.711) |  |
| **Stage 4: Bud burst** |  |  |
| **Variable** | **Coefficient (Std. Error)** |  |
| Intercept | 3.384 (1.040) | ** |
| Treatment 5 | -2.575 (1.105) | * |
| Treatment 7 | -2.837 (1.088) | ** |
| Treatment 9 | -1.576 (1.153) |  |
| Treatment 11 | -4.296 (1.099) | *** |
| City Sherbrooke | -0.104 (0.423) |  |
| **Stage 7: Full leaf expansion** |  |  |
| **Variable** | **Coefficient (Std. Error)** |  |
| Intercept | 0.672 (0.464) |  |
| Treatment 5 | 0.133 (0.669) |  |
| Treatment 7 | 0.515 (0.698) |  |
| Treatment 9 | -0.115 (0.597) |  |
| Treatment 11 | -1.905 (0.930) | * |
| City Sherbrooke | 0.153 (0.466) |  |

Cuttings that were subject to no thaws (control: treatment 3) and to one-day thaws, twice a month (treatment 7) comparatively reached stages 1 and 4 a few days earlier than other treatments (Fig. 3). Spring warming (i.e. city treatment) explained variation in phenology among cuttings at all stages, with cuttings exposed to the warmer air temperature of Montreal reaching phenological stages earlier than in Sherbrooke (Table 3, Figs. 3, 4, 5).

Overall, growing degree days accumulated by each cutting based on their thawing treatment and spring warming treatment significantly improved the explanatory power of our bud phenology models for each phenological stage tested since the AIC of the more complex model (including GDD effect) decreased by 474, 357 and 279 AIC units for the onset of budbreak (stage 1), budbreak (stage 4) and leaf fully expanded (stage 7) compared to the model only considering the effect of the thaw treatments and the average temperature difference between cities (Table 3).

## 4.0 Discussion

In this study, we integrate both climatic simulations and manipulative experiments to investigate the timely but often overlooked impact of winter thaw events on tree phenology under different scenarios of climate change. Using 12 climate models and two socioeconomic pathways (SSP-2.46 and SSP. 5.85), we first show that winter thaws are projected to increase in frequency under ongoing climate warming. We then use a manipulative experiment to determine how the leaf phenology from branch cuttings from two important tree species (sugar maple, and yellow birch) of the northern temperate forest of Eastern Canada respond to different frequencies and duration of winter thaws, and two levels of spring warming. We reveal the presence of a tipping point in the impact of these variables on leaf phenology, where cuttings exposed to the highest frequency and longest duration of thaws - in line with our climate change projections - exhibit significantly more bud mortality and later phenological development than control cuttings. We further highlight the potential for a synergistic effect leading to bud mortality under an increase in winter thaws and an increase in spring warming, underlining the importance of untangling the different drivers of tree phenology response to predict the response of forests to ongoing climate change.

### 4.1) Temporal change in projected thaw events under climate change

As expected, the ongoing climate warming is projected to increase the frequency of daily winter thaws. Contrasting with our prediction, the average duration of a winter thaw event is expected to increase marginally in the future but the occurrence of long term winter thaws in January, when the minimum air temperature remains above zero for more than one day is projected to increase from an average of three years per decade to up to eight years per decade by 2100. It is also projected to occur more frequently in December and March compared to January and February, which is expected given December and March are already warmer months compared to January and February. Therefore, future winters are expected to have more frequent days with zero-crossing and a few more episodes of prolonged thaw events, which represent a strong signal of climate change. These results are in line with the positive trends in the number of days with zero crossing projected by Nilsen et al. (2021) for inland Norway sites. Another study in southern Québec predicted that the number of zero-crossing days projected annually would decrease with climate change, despite a projected (though non-significant) increase in the number of zero-crossing days in the spring. In conjunction with our own study on projected zero-crossing days in the winter, these results suggest that the prevalence of days with zero-crossing may be expected to decrease enough during fall to counter the increase in days with zero-crossing during the winter and spring, thereby leading to a shift in the seasonal occurrence of daily thaws during the year. Initial climatic differences among our study sites and theirs could however limit comparability of our results.

We also noted inter-model variations in projected winter thaws and lack of consistency between climate models, meaning that climate models projecting the strongest and smallest trends per month varied. Moreover, we highlight that the climate model ACCESS-CM2 and ACCESS-ESM1-5 produced strongest and lowest temporal trends under the socioeconomic pathway 2.46 during the months of January and February. Therefore, when paired with different regional climate models the *Australian Community Climate and Earth System Simulator* provided both strongest and smallest temporal trends in thaw events showing the importance regional climate models have in the dynamic downscaling of the weather data such as winter thaws. Moreover, the inter-climate model variability is not yet assessed for the different climate variables and could vary in importance, likely being stronger for extreme and rare climatic events such as winter thaws compared to aggregated variables such as the mean annual air temperature. Therefore, we recommend using many pairs of global and regional climate models when assessing risk and biological responses to climate change and the inter-climate model variability is not yet assessed for each climate variables

Concerning tree physiology, the increase in thaw events in December could favor the accumulation of chilling units thereby advancing dormancy release and potentially contributing to advancing budburst. However, the increased forcing during fall under climate change could postpone the process of cold acclimation if air temperature is high enough to keep promoting growth instead of promoting chilling accumulation. Under this hypothesis, the lack of chilling could postpone bud break. On the other hand, increasing air temperature during winter could promote chilling accumulation during months when air temperature was usually well below freezing and did not usually contribute to chilling accumulation. Hence, chilling accumulation could shift from October to December without impacting the amount of accumulated chilling. Moreover, the increase in daily thaws during the months of January, February and March could contribute in adding forcing that would not be considered by the growing degree-day calculation, hence increasing the discrepancy between the real accumulated forcing by trees and the accumulated forcing considered by the growing degree day calculation. Therefore, the difference between the real timing of bud break and the timing of bud break projected by the ecophysiology models might increase under climate change, adding a source of uncertainty to bud break phenology predictions. While calculating hourly growing degree days instead of daily growing degree-days would help alleviate this issue [44, 45], most climate models, such as the ones we obtained climate data from, only provide the daily maximum and minimum air temperature from which the mean daily air temperature is calculated because storing and sharing these data is less computationally intensive. With increasing technology and compute power, the hourly data could become increasingly useful and important to understand the future winter thaw regime and its impact on tree phenology and growth. However, there is an important dilemma when using hourly, even daily data from climate models which is that models are simplified representations of our understanding of the Earth processes, as such, models are relevant to identify long term trends such as differences between 30 year periods but are not built to predict accurate short-term events such as winter thaws [46]. Hence, the use of hourly data from climate models could prove advantageous to ecophysiological models that run a short time scale (hourly or daily) but could also prove disadvantageous if the weather projections are not accurate [44, 45].

Since photoperiod is a strong environmental cue triggering growth cessation and cold acclimation [21], we expect that the future frequency and duration of thaw events will impact the accumulation of chilling and forcing so that it will be decoupled from the photoperiod signal and could potentially lead to novel timing of tree phenology or increase the variability in bud break timing, hence, reducing our ability at predicting it.

### 4.2) Effects of winter thaws and spring warming on bud phenology

As shown using climate projections, the frequency and intensity of winter thaws could increase as a result of climate change, increasing the divergence between the real and predicted state of cold hardiness, chilling and forcing in trees and potentially impacting their vegetative phenology. Using a manipulative experiment, we first tested the hypothesis that increasing the number and duration of winter thaws would advance the onset of the bursting of buds by either contributing to the chilling needs or the forcing in spring. We predicted that the timing of bud burst of tree cuttings exposed to experimental winter thaw treatments would be correlated with the duration and frequency of the thaw events, with cuttings exposed to more thawing bursting first.

Unlike expected, we showed that winter thaws increased bud mortality and delayed bud opening for the surviving cuttings. Failure to reach bud burst and delays in phenological development were most extreme in the thawing treatment with the longest and most frequent winter thaws, revealing a tipping point in the impact of winter thaws on yellow birch bud mortality and phenology. While cuttings unable to open their buds were more numerous in most treatments than in the control, except for the most extreme treatments, dates at which branches reached bud burst or full leaf expansion were similar to those of control branches. Rather than allowing the trees to open their leaves earlier and potentially benefit from a longer growing season, these results suggest that increased levels of thawing in the winter could negatively impact the volume of the canopy by decreasing the proportion of buds able to develop, and thus incur energetic costs for the trees in reducing the leaf surface available for photosynthesis, or in producing new leaves from dormant buds. The inability of sugar maple to reactivate its buds could suggest that this species required more chilling than we provided or that the lack of chilling delayed bud break timing. Providing only water to the cuttings could also have been insufficient for triggering the bursting of buds if plant hormones were to play a larger role for that species than for yellow birch. This experiment shows that different physiological processes may be required for bud reactivation among the two species.

By exposing the cuttings to two different temperature regimes in the spring, we further tested the hypothesis that increasing the forcing in spring would advance the onset of the bursting of buds irrespective of the thaw treatment. As predicted, cuttings at the warmer site opened their buds earlier compared to the colder site, an effect that can be explained through differential accumulation of growing degree days among sites. These results suggest that unlike degree-days accumulated during the winter that can lead to an increased mortality of cuttings in extreme cases, those accumulated in the spring contribute to accelerating bud development. Our study therefore adds to the large body of literature showing the advance of bud phenology under increasing spring forcing [22, 47, 48] but is particularly novel in showing the effects of winter thaw on bud mortality and phenological delays. In accordance with our results, warm winters were also found to delay spring phenology on the Tibetan Plateau [49] but had the opposite effect in Europe [50]. The latter study however provided longer and more constant (i.e. greenhouse-controlled) warming to the buds than in our own experiment, suggesting that pulsed thaw events of shorter durations but higher frequency as simulated in our experiment may have more negative impacts on bud development than the mean increase in air temperature of warmer winters. Given Europe is warmer than Québec and the Tibetan Plateau, tree species might also respond differently to warming when originating from a cold or a warm environment. We acknowledge that in both cities, bud mortality could have been promoted by sub-zero temperatures happening during the spring. While sub-zero temperatures were detected in Sherbrooke during the spring, bud mortality was higher in Montreal where no such event was detected for all the time that cuttings were outside, such that bud mortality in our experiment is mostly associated with thaw treatments instead of being associated with spring freezing.

Overall, the phenology and survival of buds in the face of climate change could possibly depend on a synergistic effect between the frequency and duration of thaw treatments and the level of spring warming, with high frequency and long duration of thaw events leading to increased bud mortality under higher spring warming. Indeed, bud mortality was higher in Montreal compared to Sherbrooke, and only the spring forcing varied between cities. Therefore, winter forcing linked with thaw events as well as spring forcing could increase bud mortality compared to warming in winter only, but this hypothesis remains to be tested.

Even if progress in determining the physiological mechanisms regulating sensitivity to photoperiod and temperature is ongoing [21, 25], process-based models predicting tree phenology such as the unified model of budburst which require the parametrization of nine parameters, are currently only based on air temperature [16, 27]. We show that taking into account the frequency and duration of winter thaws is crucial in developing more precise models of bud phenological development in the face of climate change. Recent studies also showed that photoperiod, precipitation and the microbiome also played a role in controlling bud phenology [51, 52, 53] which could importantly increase the complexity of ecophysiology models and continue to improve our capacity to predict changes in tree phenology under climate change.

## Conclusion

We experimentally tested the effect of winter thaws on tree phenology of two important tree species of the northern temperate American forests. We used a novel yet simple method to mimic the effect of winter thaws by keeping cuttings in freezers and periodically leaving them out to + 4 °C for different durations and frequencies. While exposure to short durations (one day) and low frequencies (once a month) did not significantly advance yellow birch’s bud phenology compared to the control (no thaw), frequent (twice a month) and long (three days) exposure to winter thaws delayed bud break timing and increased bud mortality. Since climate models project an increase in daily winter thaws, we could expect future bud break timing to become earlier, except during the years when thaw events of long durations will occur, which could delay the timing of budbreak and induce mortality of buds. In such a case, we could expect less foliage in forest canopy, at least until new leaves will be formed, which could impact ecosystem services provided by the northern temperate forest and more specifically yellow birch such as carbon sequestration. Overall, our experiment revealed important differences in tree physiology between sugar maple and yellow birch and showed that by increasing bud mortality, winter thaws could impact tree phenology even if these events are rare and of short durations they should be better considered by global vegetation models and ecophysiological models predicting future bud break timing. We also highlighted inter-climate model variability in winter thaw projections, hence assessing risk under climate change starts by using many different pairs of global and regional climate models during the assessment.

## Acknowledgements

We are thankful to the members of the National Parc of Mont-Orford to have granted us access to the site.

## Author contributions

Both co-authors contributed equally at every step of the research project.

B.M.: Conceptualization, Data, Analyses, Writing

DouceG.L.: Conceptualization, Data, Analyses, Writing

## Conflict of interest

No conflict of interest declared

## Funding

This research did not receive any specific grant from funding agencies in the public, commercial, or not-for-profit sectors.

## Data availability

The phenology data as well as the air temperature data collected by the authors will be deposited on Figshare upon publication/acceptance of the manuscript. The projected daily climate data analyzed in this article is already available from the Pacific Climate Impacts Consortium (PAVICS) data portal (https://data.pacificclimate.org/portal/downscaled_cmip6/map/).

## Notes

### Competing Interest Statement

The authors have declared no competing interest.

