## Supplementary Material for "Experimental exposure to winter thaws reveals tipping point in yellow birch bud mortality and phenology in the northern temperate forest of Québec, Canada"

### Supplementary Information

**Table S1**. Description of all the treatments tested to test the effect of winter thaws on the timing of budbreak under climate change in our experimental design.

| ID | Spring phenology | | | Winter thaws | | | | |
|---|---|---|---|---|---|---|---|---|
| | Field | Warm (Montreal) | Cold (Sherbrooke) | Control | Intensity (4 °C) | | Frequency | |
| | | | | No exposure to thaw events | Low (24 hours) | High (72 hours) | Low (once a month) | High (twice per month) |
| 1 | Yes | - | - | - | - | - | - | - |
| 2 | Yes | - | - | - | - | - | - | - |
| 3 | - | Yes | - | Yes | - | - | - | - |
| 4 | - | - | Yes | Yes | - | - | - | - |
| 5 | - | Yes | - | - | Yes | - | - | - |
| 6 | - | Yes | - | - | - | Yes | - | - |
| 7 | - | Yes | - | - | - | - | Yes | - |
| 8 | - | Yes | - | - | - | - | - | Yes |
| 9 | - | - | Yes | - | Yes | - | - | - |
| 10 | - | - | Yes | - | - | Yes | - | - |
| 11 | - | - | Yes | - | - | - | Yes | - |

| | | | | | | | | |
|---|---|---|---|---|---|---|---|---|
| 12 | - | - | Yes | - | - | - | - | Yes |

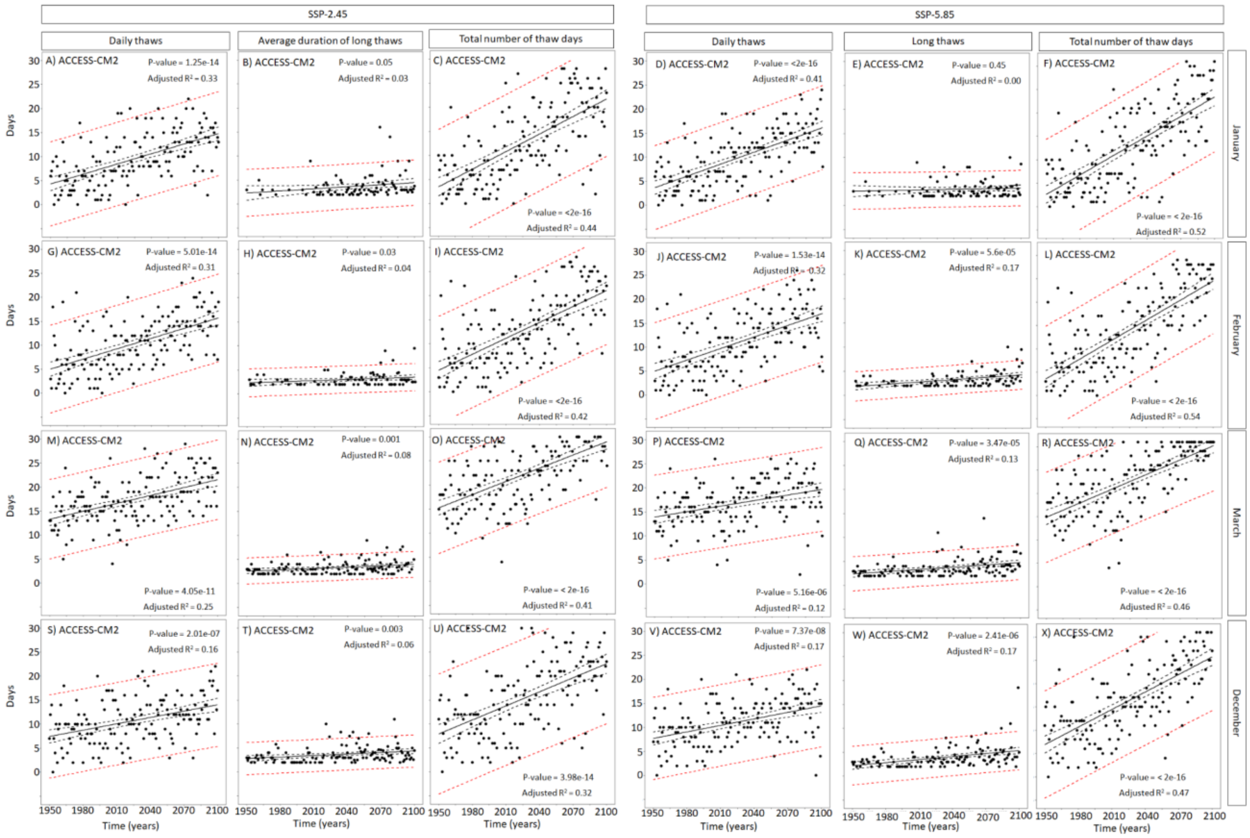


**Figure S1**. Temporal trends in the number of thaws lasting a day (daily thaws), the average duration of a thaw event and finally the total number of thaw events (daily + long term thaws) projected per winter months under two socioeconomic pathways for the climate model ACCESS-CM2. The plain black line represents the linear trend, the dashed black lines represent the 95 % confidence interval and the dashed red lines represent the prediction interval. Summary of the linear trends are reported in the figures.

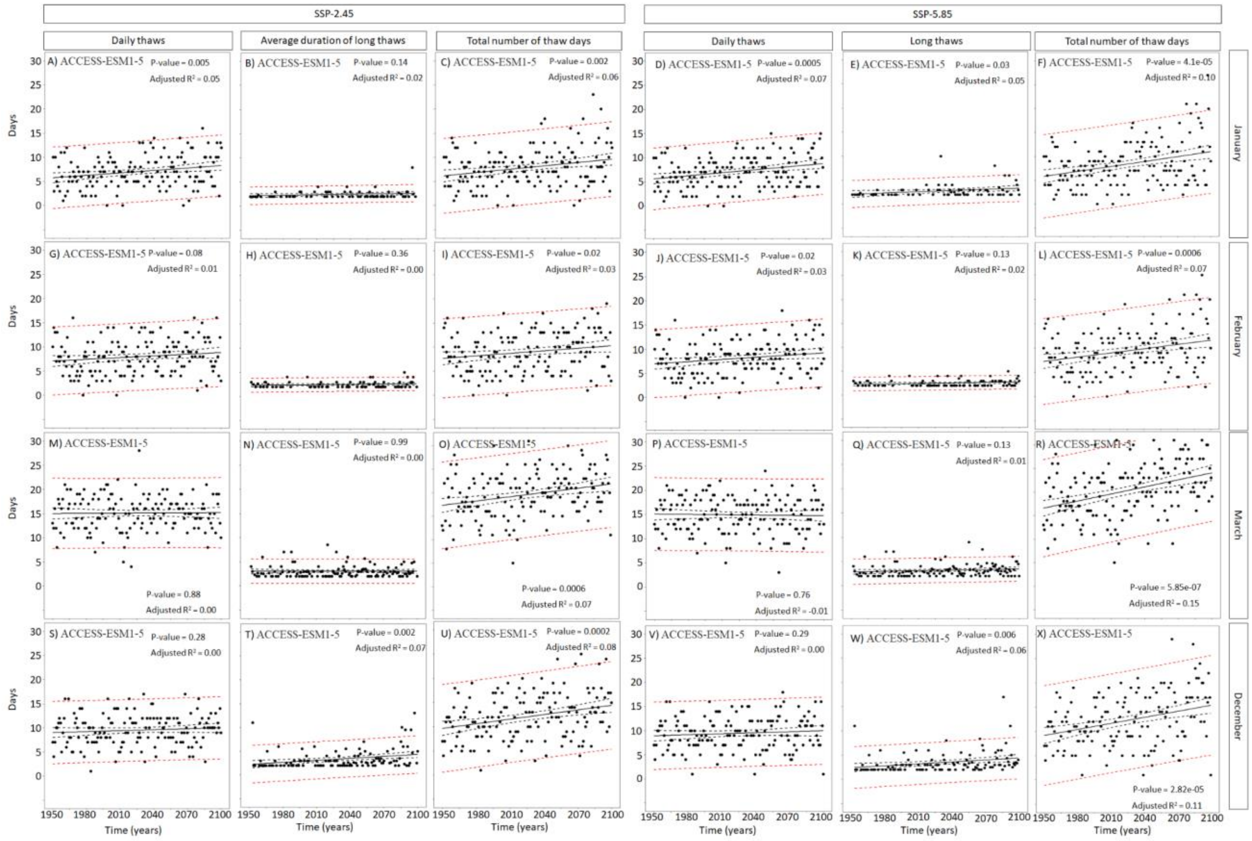

**Figure S2.** Temporal trends in the number of thaws lasting a day (daily thaws), the average duration of a thaw event and finally the total number of thaw events (daily + long term thaws) projected per winter months under two socioeconomic pathways for the climate model ACCESS-ESM1-5. The plain black line represents the linear trend, the dashed black lines represent the 95 % confidence interval and the dashed red lines represent the prediction interval. Summary of the linear trends are reported in the figures.

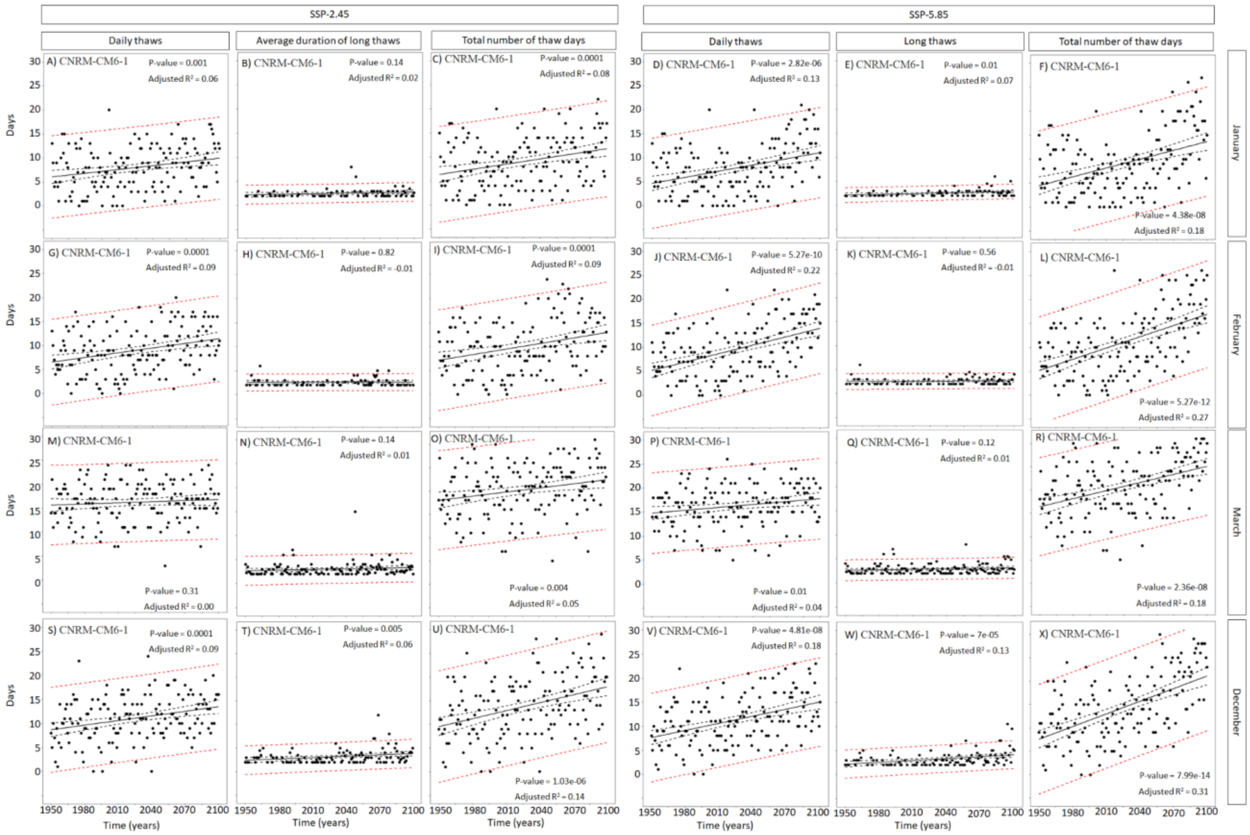

**Figure S3.** Temporal trends in the number of thaws lasting a day (daily thaws), the average duration of a thaw event and finally the total number of thaw events (daily + long term thaws) projected per winter months under two socioeconomic pathways for the climate model CNRM-CM6-1. The plain black line represents the linear trend, the dashed black lines represent the 95 % confidence interval and the dashed red lines represent the prediction interval. Summary of the linear trends are reported in the figures.

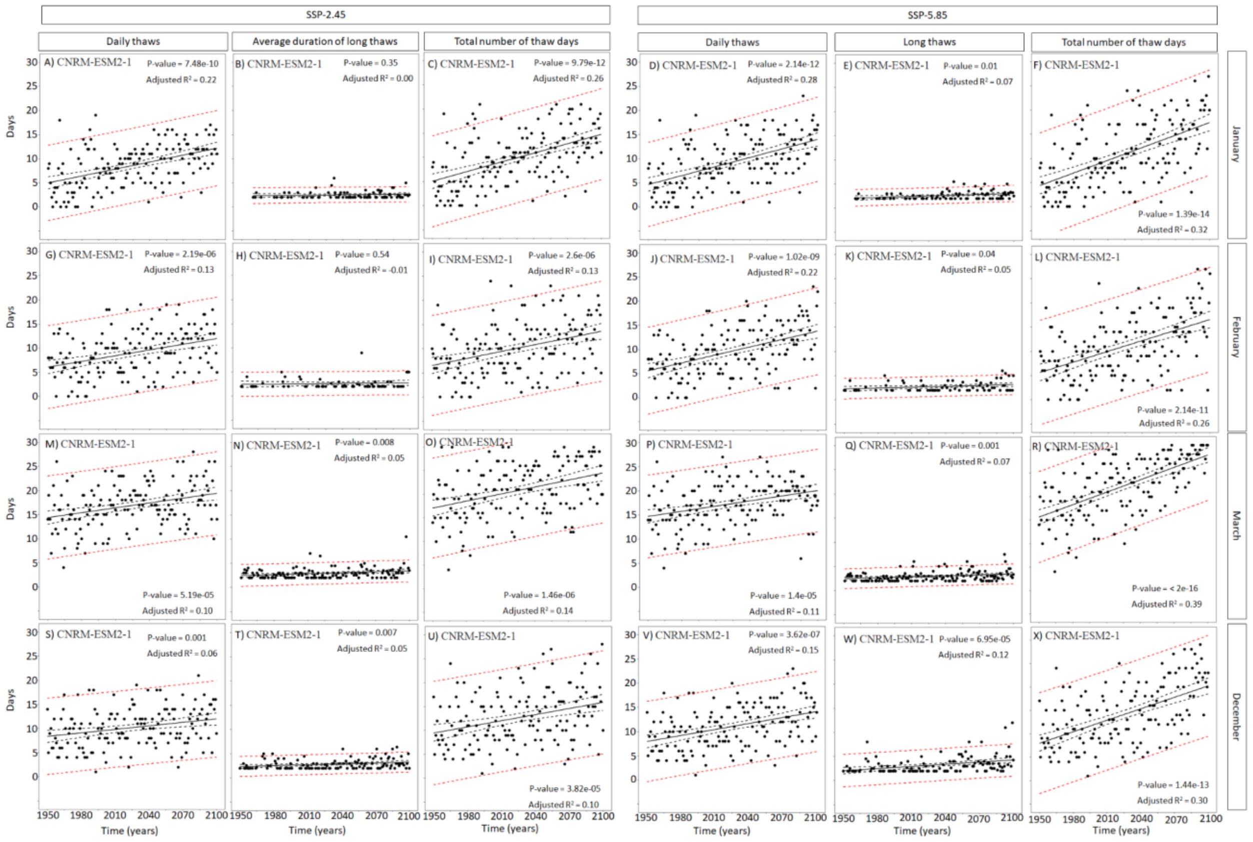


**Figure S4.** Temporal trends in the number of thaws lasting a day (daily thaws), the average duration of a thaw event and finally the total number of thaw events (daily + long term thaws) projected per winter months under two socioeconomic pathways for the climate model CNRM-ESM2-1. The plain black line represents the linear trend, the dashed black lines represent the 95 % confidence interval and the dashed red lines represent the prediction interval. Summary of the linear trends are reported in the figures.

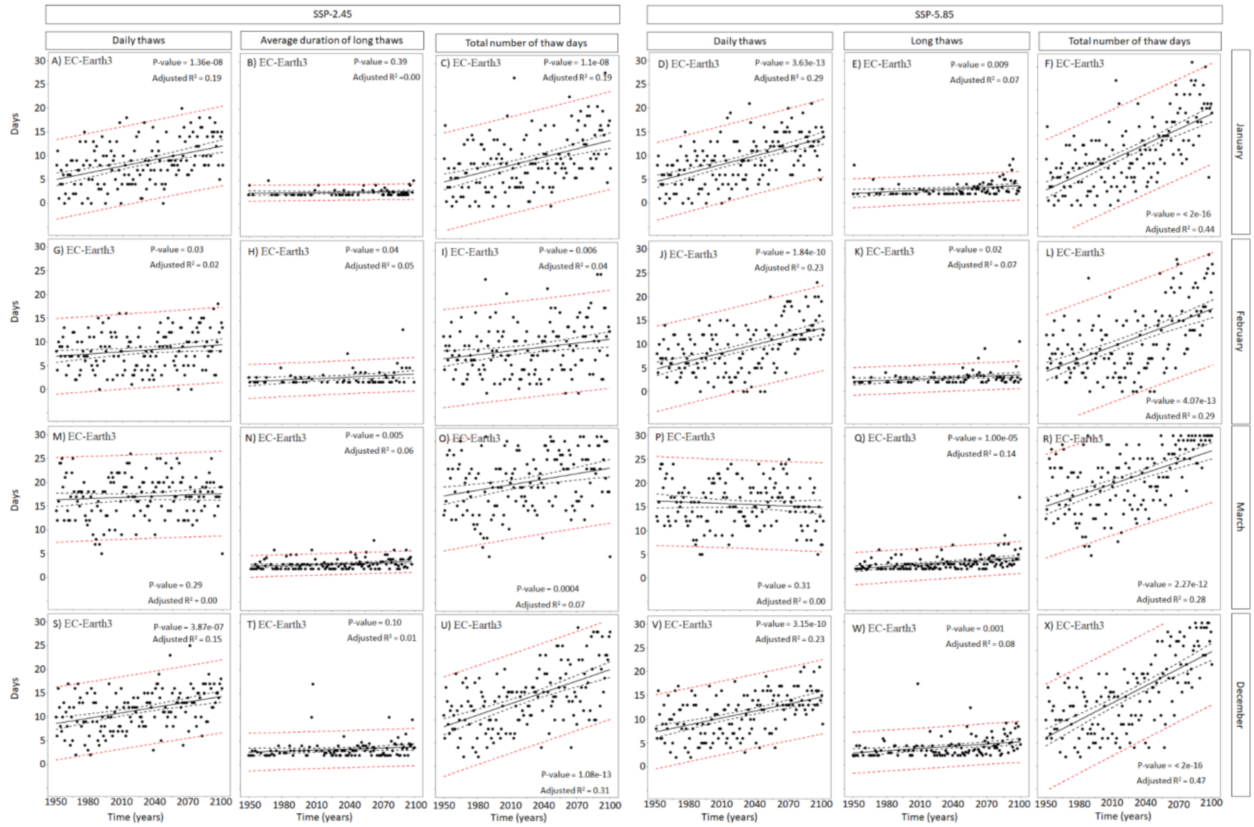

**Figure S5.** Temporal trends in the number of thaws lasting a day (daily thaws), the average duration of a thaw event and finally the total number of thaw events (daily + long term thaws) projected per winter months under two socioeconomic pathways for the climate model EC-Earth3. The plain black line represents the linear trend, the dashed black lines represent the 95 % confidence interval and the dashed red lines represent the prediction interval. Summary of the linear trends are reported in the figures.

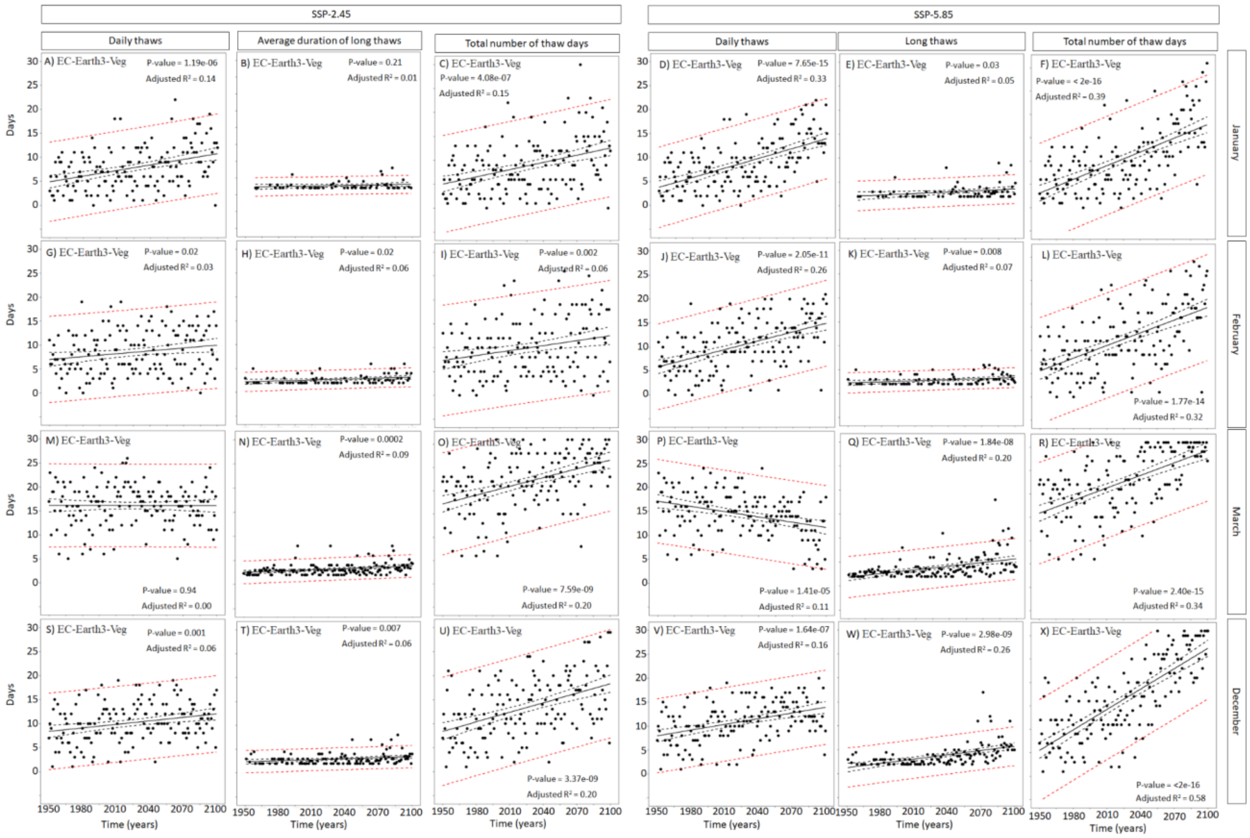

**Figure S6.** Temporal trends in the number of thaws lasting a day (daily thaws), the average duration of a thaw event and finally the total number of thaw events (daily + long term thaws) projected per winter months under two socioeconomic pathways for the climate model EC-Earth3-Veg. The plain black line represents the linear trend, the dashed black lines represent the 95 % confidence interval and the dashed red lines represent the prediction interval. Summary of the linear trends are reported in the figures.

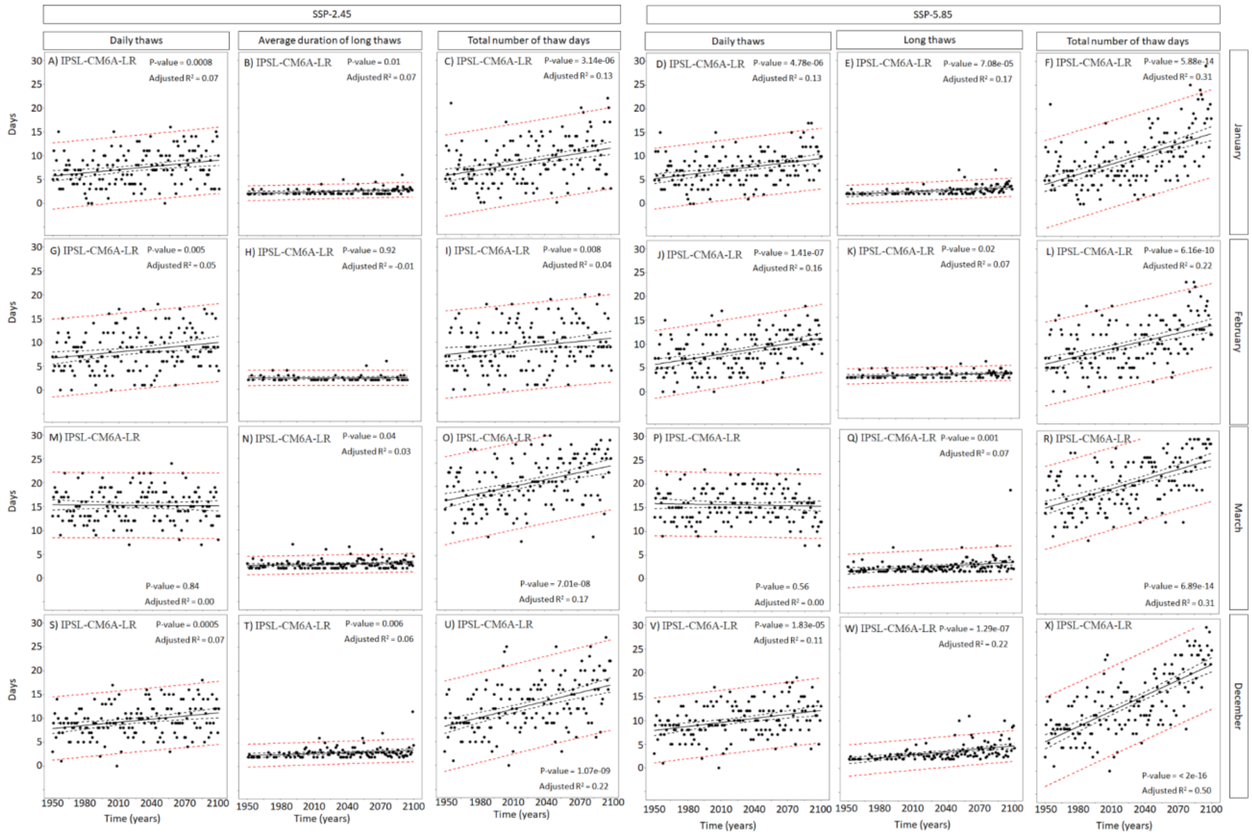


**Figure S7.** Temporal trends in the number of thaws lasting a day (daily thaws), the average duration of a thaw event and finally the total number of thaw events (daily + long term thaws) projected per winter months under two socioeconomic pathways for the climate model IPSL-CM6A-LR. The plain black line represents the linear trend, the dashed black lines represent the 95 % confidence interval and the dashed red lines represent the prediction interval. Summary of the linear trends are reported in the figures.

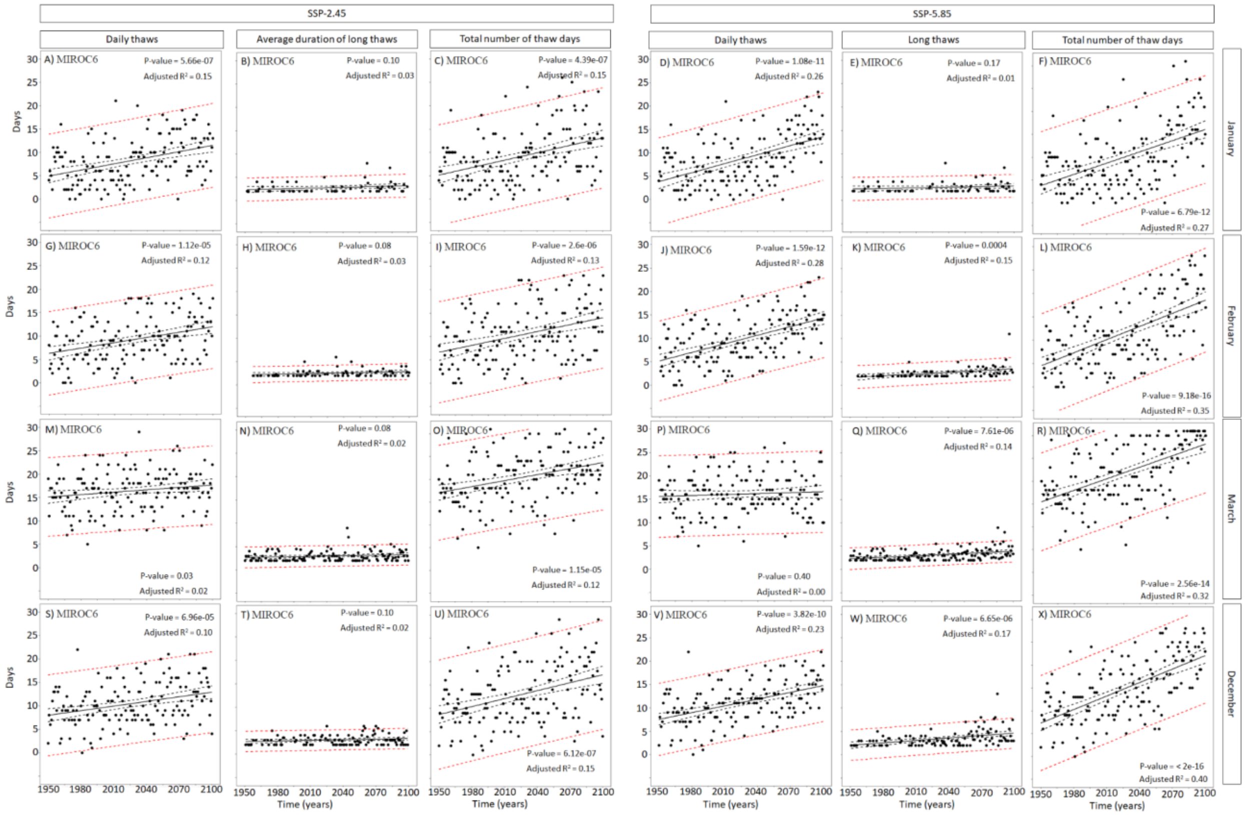


**Figure S8.** Temporal trends in the number of thaws lasting a day (daily thaws), the average duration of a thaw event and finally the total number of thaw events (daily + long term thaws) projected per winter months under two socioeconomic pathways for the climate model MIROC6. The plain black line represents the linear trend, the dashed black lines represent the 95 % confidence interval and the dashed red lines represent the prediction interval. Summary of the linear trends are reported in the figures.

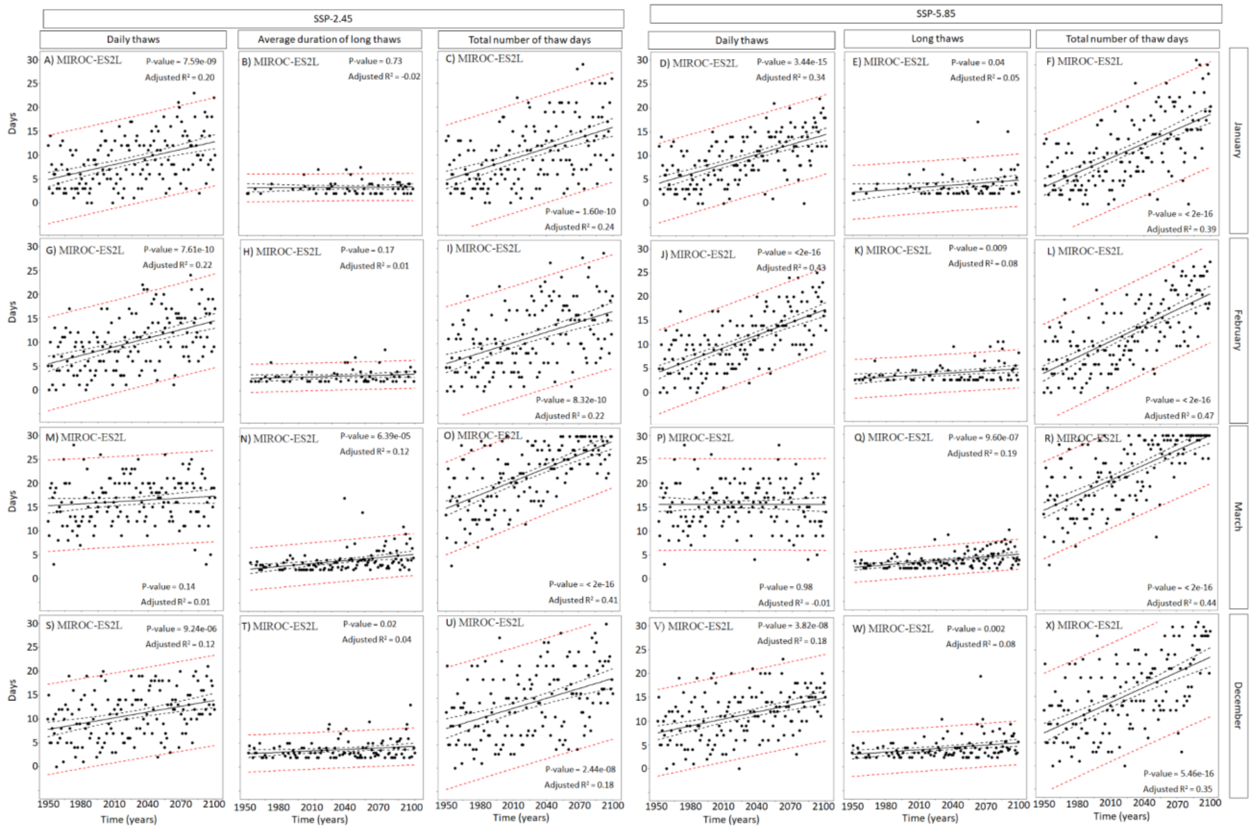

**Figure S9**. Temporal trends in the number of thaws lasting a day (daily thaws), the average duration of a thaw event and finally the total number of thaw events (daily + long term thaws) projected per winter months under two socioeconomic pathways for the climate model MIROC-ES2L. The plain black line represents the linear trend, the dashed black lines represent the 95 % confidence interval and the dashed red lines represent the prediction interval. Summary of the linear trends are reported in the figures.

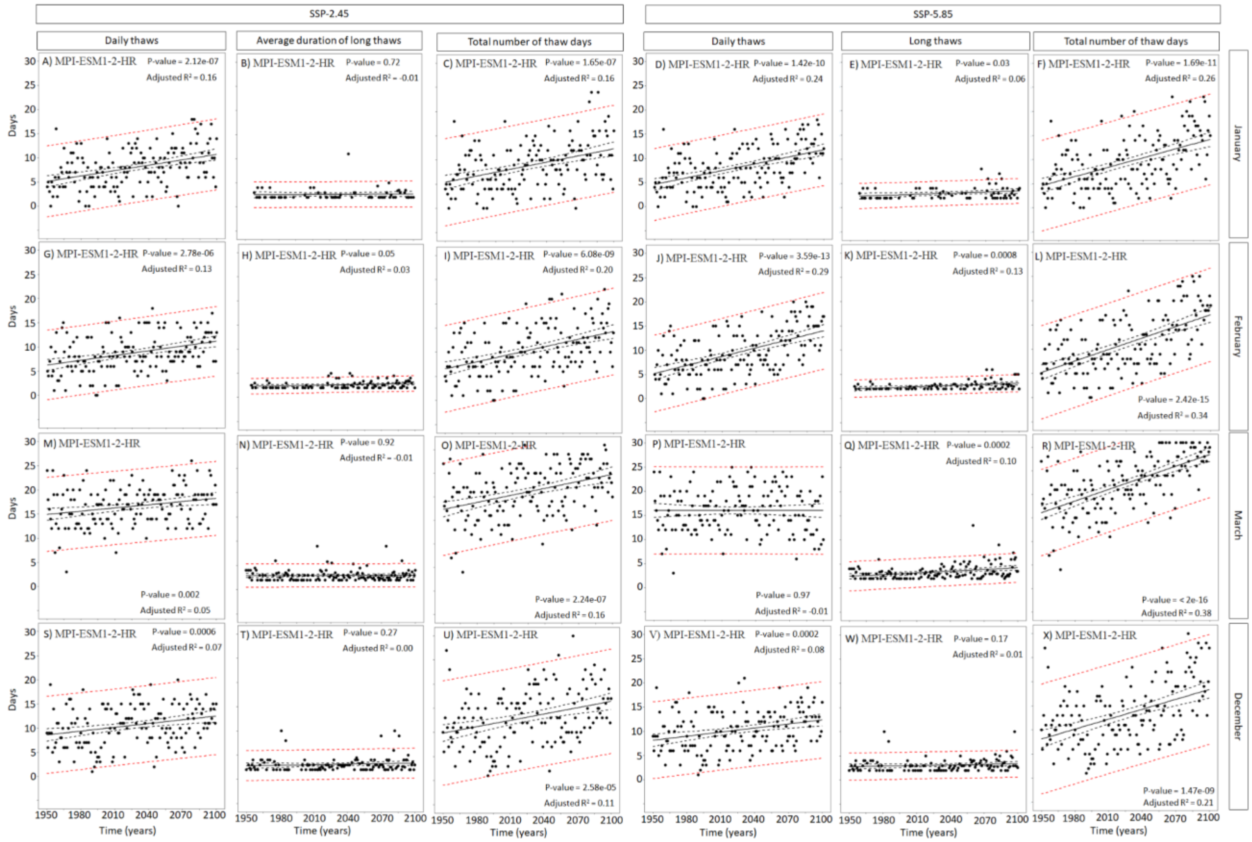

**Figure S10.** Temporal trends in the number of thaws lasting a day (daily thaws), the average duration of a thaw event and finally the total number of thaw events (daily + long term thaws) projected per winter months under two socioeconomic pathways for the climate model MPI-ESM1-2-HR. The plain black line represents the linear trend, the dashed black lines represent the 95 % confidence interval and the dashed red lines represent the prediction interval. Summary of the linear trends are reported in the figures.

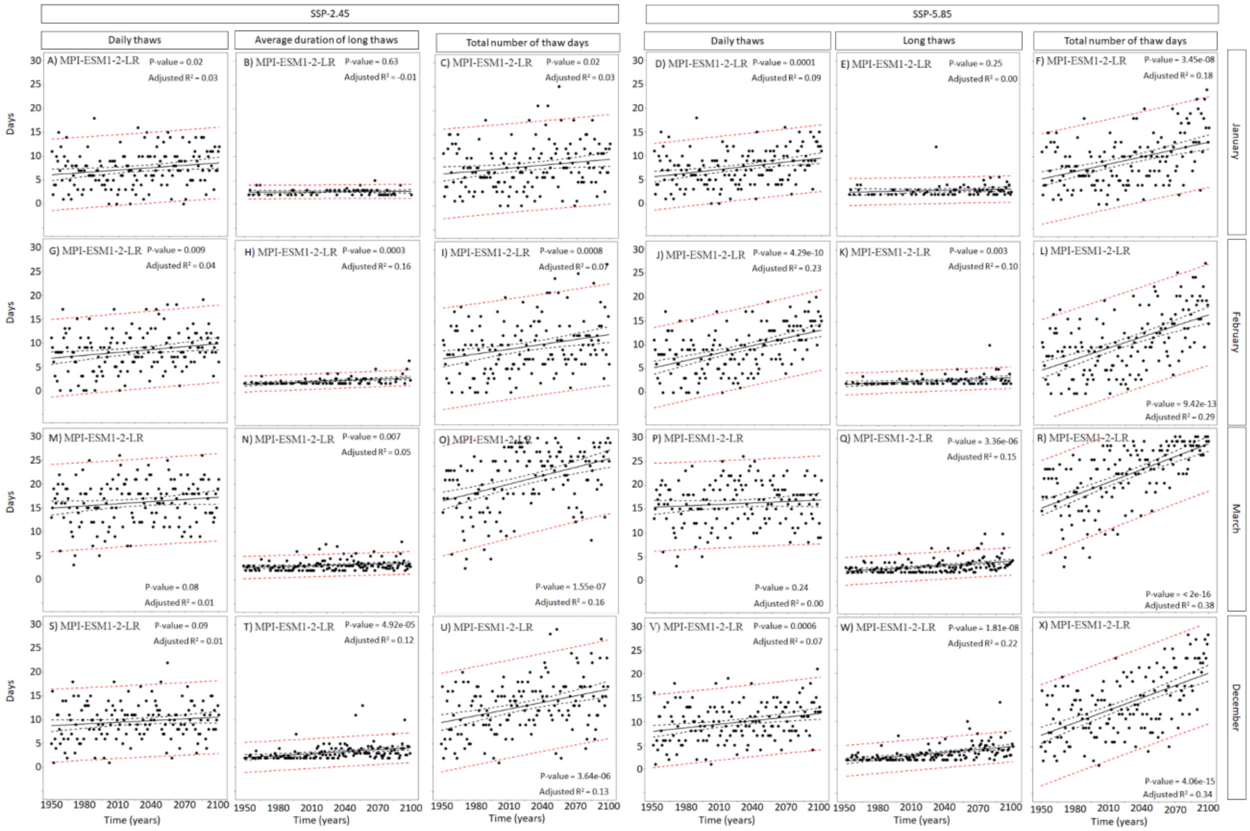


**Figure S11.** Temporal trends in the number of thaws lasting a day (daily thaws), the average duration of a thaw event and finally the total number of thaw events (daily + long term thaws) projected per winter months under two socioeconomic pathways for the climate model MPI-ESM1-2-LR. The plain black line represents the linear trend, the dashed black lines represent the 95 % confidence interval and the dashed red lines represent the prediction interval. Summary of the linear trends are reported in the figures.

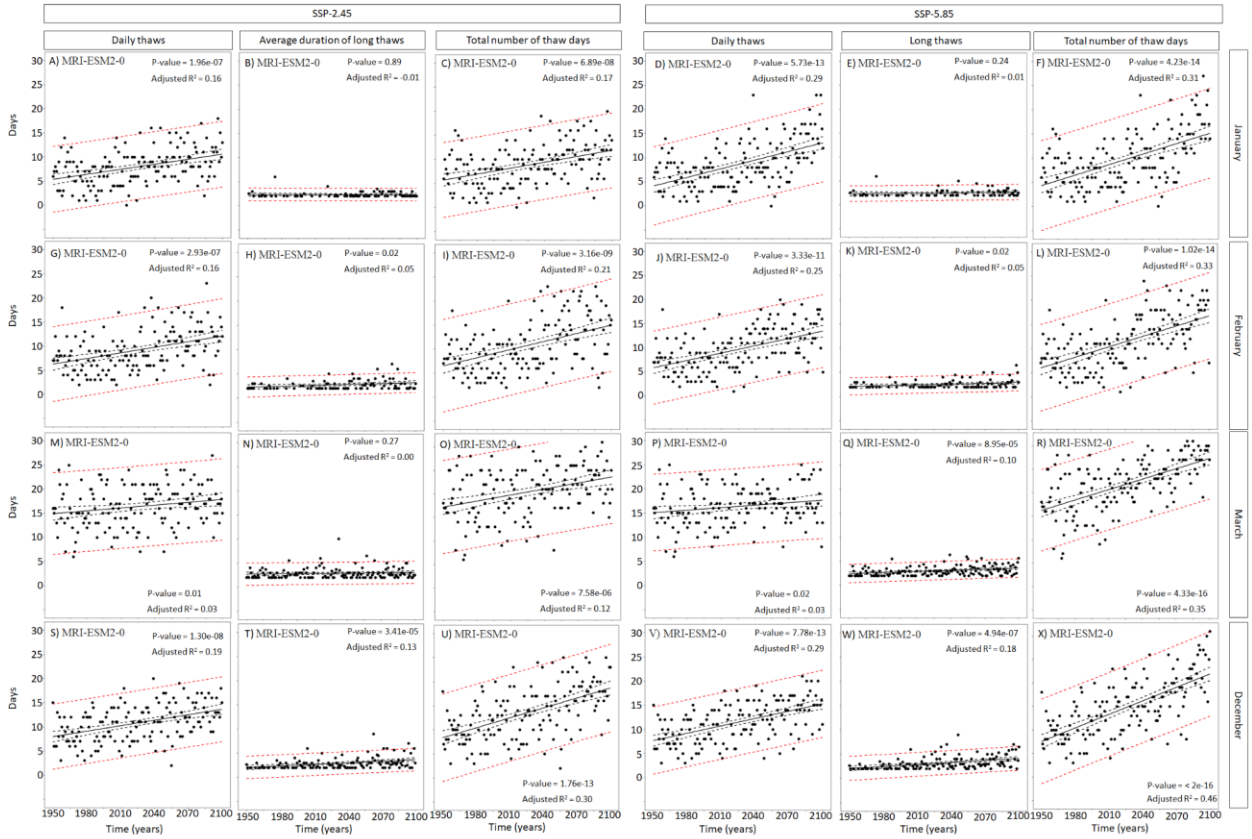

**Figure S12.** Temporal trends in the number of thaws lasting a day (daily thaws), the average duration of a thaw event and finally the total number of thaw events (daily + long term thaws) projected per winter months under two socioeconomic pathways for the climate model MRI-ESM2-0. The plain black line represents the linear trend, the dashed black lines represent the 95 % confidence interval and the dashed red lines represent the prediction interval. Summary of the linear trends are reported in the figures.

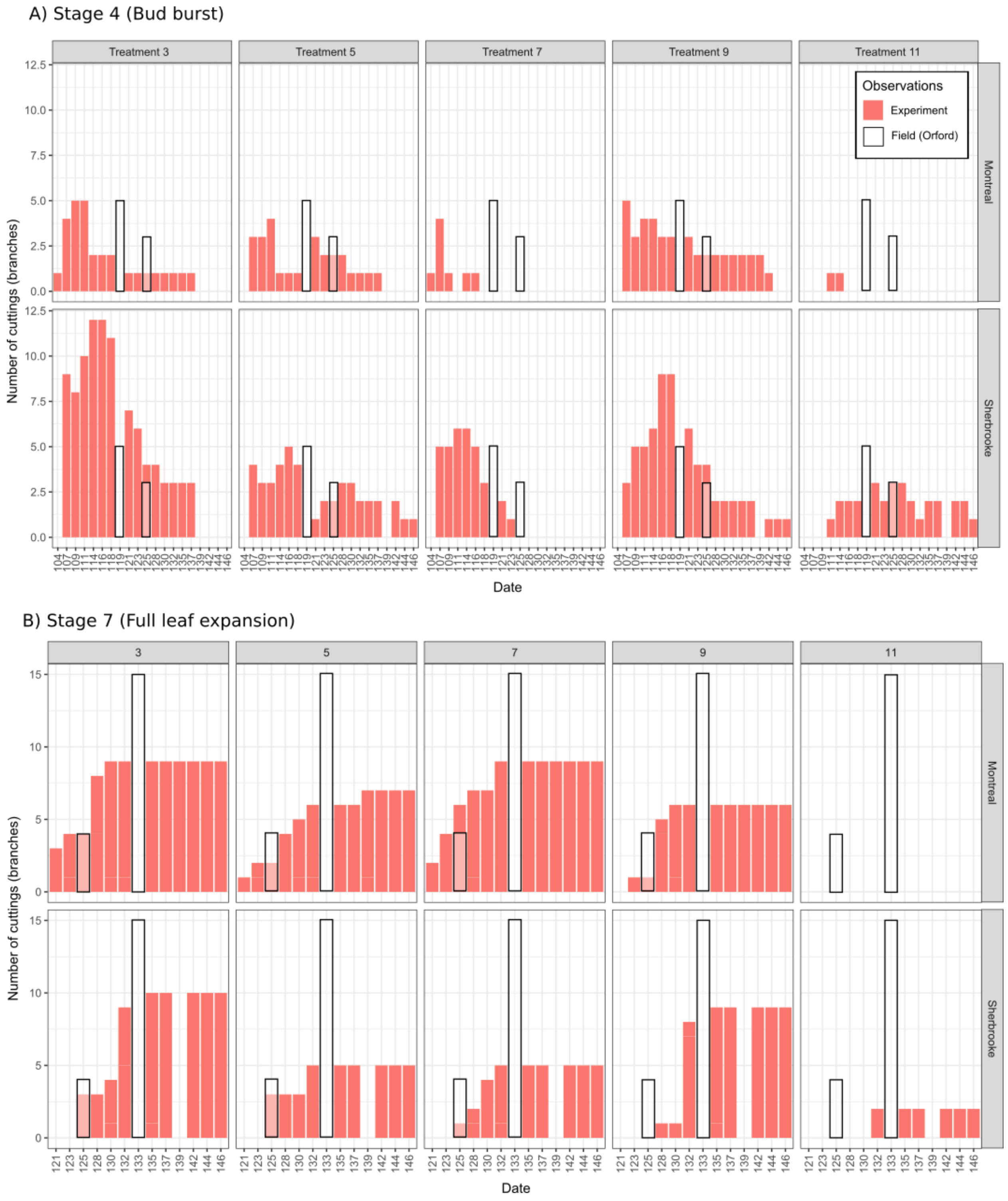


**Fig. S13.** Number of cuttings reaching phenological stages 4 (panel A) and 7 (panel B) in the experiment (red bars) relative to phenological stages of branches from the same trees observed in the field (black border bars). Data presented for yellow birch alone.

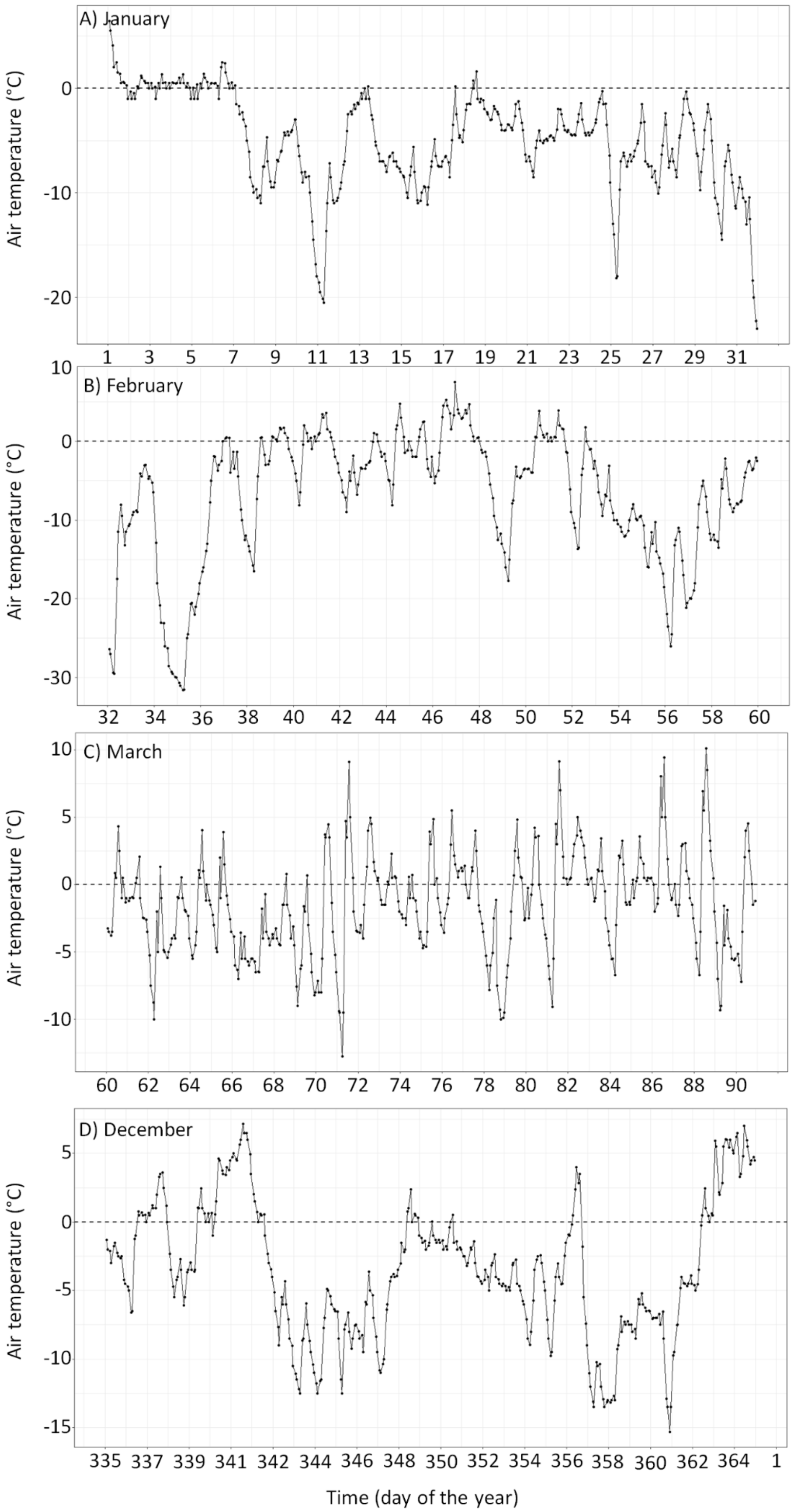


**Fig. S14.** Air temperature averaged over the seven air temperature loggers set *in situ* at the Mont-Orford National Park during the winter months.

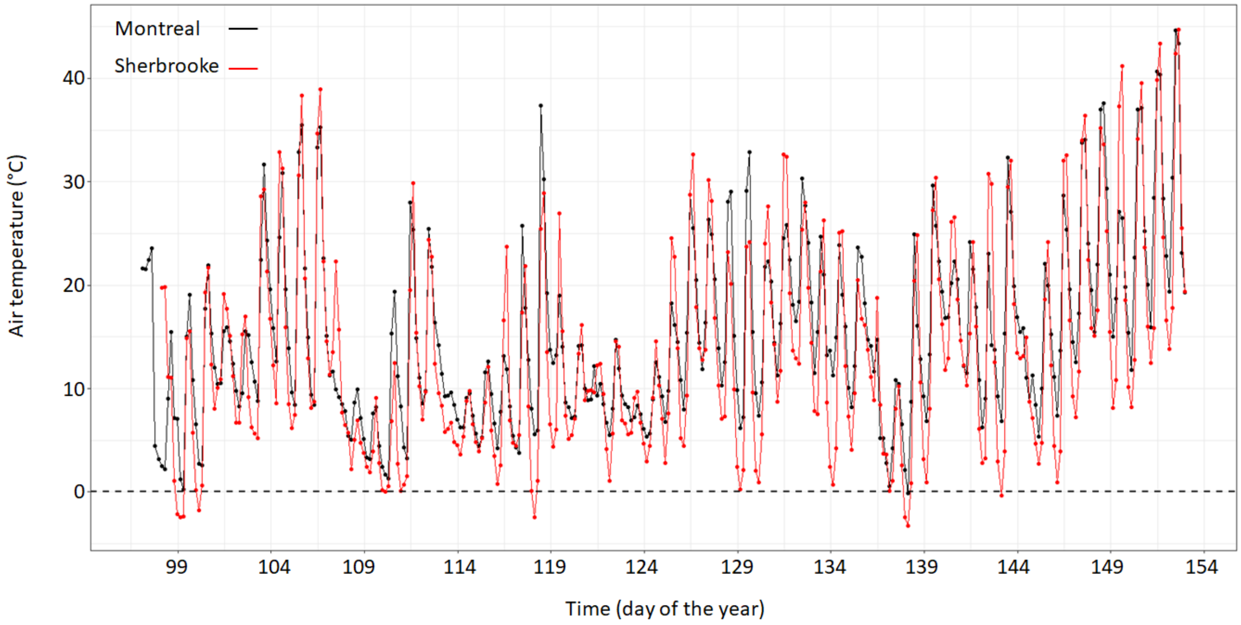


**Fig. S15.** Hourly average air temperature of all the iButtons that were set at each city (Montreal and Sherbrooke) during spring 2023 for the regrowth test.

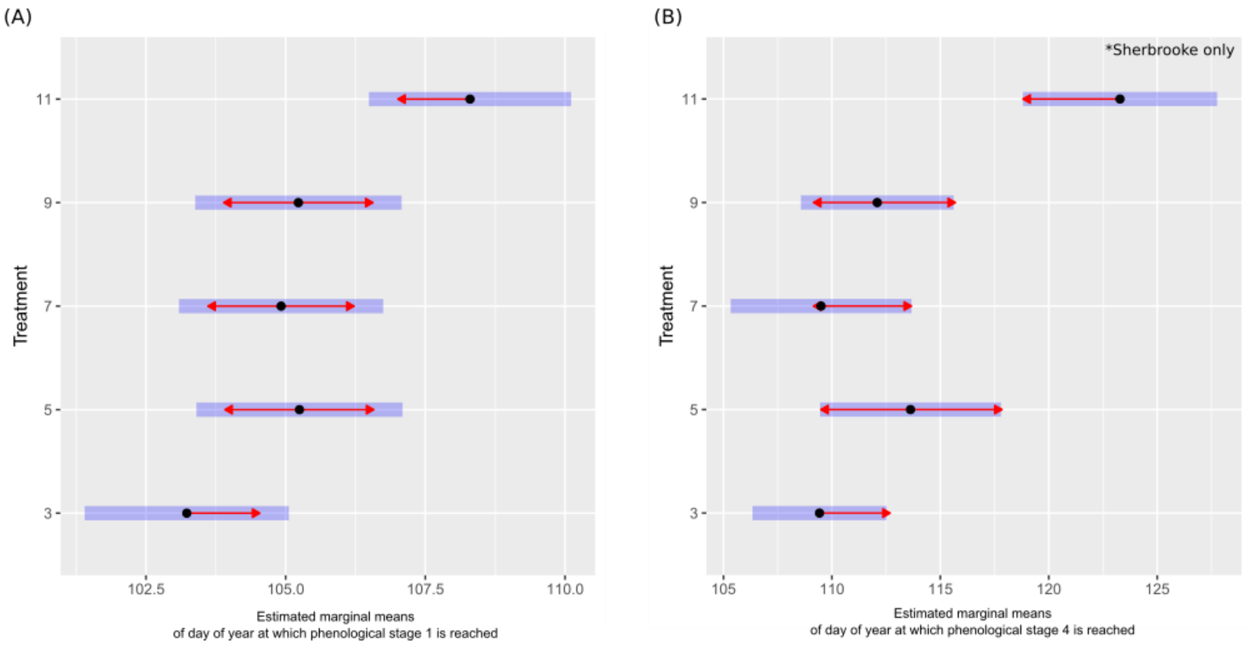


**Fig. S16.** Effect of treatment on day of year at which stage 1 (Panel A) and stage 4 (Panel B) phenological stages are reached. Panel B presents data for Sherbrooke only because not enough observations could be made for Montreal. Black dots represent the value of estimated marginal means of day of year at which these stages are reached. The blue bars are confidence intervals. The absence of overlap among red arrows of two treatments indicates that the mean day of year at which cuttings reach that stage are significantly different among these treatments. These pairwise comparisons were adjusted for multiple testing using a Tukey adjustment.
